# Convergent evolution of sex chromosomes in two palm species, *Phoenix dactylifera* and *Kerriodoxa elegans*

**DOI:** 10.1101/2024.06.27.600560

**Authors:** H. Tessarotto, T. Beulé, E. Cherif, J. Orjuela, P. Farhat, A. Lindström, A. Lemansour, S. Santoni, J. Käfer, F. Aberlenc

## Abstract

- In palms, many dioecious species have emerged from at least 8 independent events, yet the mechanisms of sex determination remain poorly understood. Here, we identify and compare the sex chromosomes of *Kerriodoxa elegans* with those of the well-studied date palm (*Phoenix dactylifera*), which evolved dioecy independently from a hermaphroditic common ancestor.
- We developed target sequence capture kits and inferred sex-linked genes using a probabilistic approach in both species.
- We find a striking similarity between the sex-linked regions of *K. elegans* and *P. dactylifera*, with the majority of sex-linked genes being common between the two species. However, we confirm that these regions evolved independently, much later than the split between the lineages.
- This case of convergent evolution seems to be unique in plants so far, and raises questions on the mechanisms of sex determination. A possible explanation is that this region contains genes related to flower sexual development, and is syntenic in monoecious palm species as well, suggesting it might serve as a genetic ‘toolbox’ for flower unisexualization in palms.

## Introduction

Flowering plants are characterized by a large diversity in sexual systems. While most species have bisexual flowers (the presumed ancestral state, Sauquet *et al*., 2017), unisexual flowers evolved rather frequently in many different clades. About 10% of species have unisexual flowers (Renner, 2014), either with flowers of both sexes on one plant (monoecy) or on different individuals (dioecy). Of about 15,000 dioecious angiosperm species, sex determination is only known in fewer than 100, distributed in about 30 genera (Leite Montalvão *et al*., 2021; Charlesworth D & Harkess A, 2024). In most of these species, sex chromosomes are XY (male heterogamety), with small non-recombining regions, although there are some notable exceptions (Renner & Müller, 2021).

Sex chromosomes evolve from a pair of autosomes and once they have stopped recombining, the sex-limited chromosome (Y in XY systems) degenerates. This has dramatic consequences: deleterious mutations accumulate and many genes cease to function and can be lost (Ming *et al*., 2011). However, what causes the initial recombination suppression and subsequent extensions of this region, leading to striking dissimilarities among species, remains poorly understood (Renner & Müller, 2021). In plants, sex-determining genes probably evolved independently with each transition to unisexuality, implying that they must be identified from scratch in each clade, and so far only a few such genes have been characterized (Henry *et al*., 2018; Leite Montalvão *et al*., 2021; Renner & Müller, 2021). How sex-determining genes might give rise to the initiation and extension of non-recombining regions typical of sex chromosomes remains an open question (Charlesworth & Charlesworth, 1978; Ponnikas *et al*., 2018; Renner & Müller, 2021). Recent modeling studies have identified new possible mechanisms (Jeffries *et al*., 2021; Lenormand & Roze, 2022; Jay *et al*., 2022) but their predictions remain to be tested (Käfer, 2022).

The study of plants, and their repeated evolution of dioecy and sex chromosomes, could shed light on the generality of the evolutionary patterns and the processes behind them. In a few groups, comparative analyses of related species have been performed, yielding different results in each case. For example, in the genus *Silene*, that contains the model species *S. latifolia*, dioecy evolved independently in at least three groups (Marais *et al*., 2011). In two of these, species with sex chromosomes have been identified, with one group consisting of five species that share an ancestral well-differentiated sex chromosome pair, while in the other group sex-linked regions are small and likely to have undergone turnovers (Balounova *et al*., 2019). In the Salicaceae family, dioecy is ancestral but different sex chromosome pairs have been found in different species, once again suggesting sex chromosome turnover (Hu N *et al*., 2023). The two sister genera *Cannabis* and *Humulus* inherited a homologous sex chromosome system from their common ancestor with subsequent expansion of the non-recombining region in both, thus leading to a rather unique situation in plants during which quite degenerated sex chromosomes with a large non-recombining region have remained stable for more than 20 million years (Prentout *et al*., 2021). The reasons for these differences are not understood.

Arecaceae (palms) are a large family of flowering plants with about 2,600 species (‘PALMweb’, 2025). It has a high proportion of species with unisexual flowers, with 52% being monoecious and 30% dioecious, as well as many transitions between these sexual systems. Consequently, it has been called “a microcosm of sexual system evolution” (Nadot *et al*., 2016). Ancestral state reconstructions do not provide a consensus on whether the common ancestor of all palms exhibited monoecy or hermaphroditism, but it is clear that dioecy has evolved independently in several palm clades (Figure 1; Nadot *et al*., 2016; Cássia-Silva *et al*., 2021).

**Figure 1:**
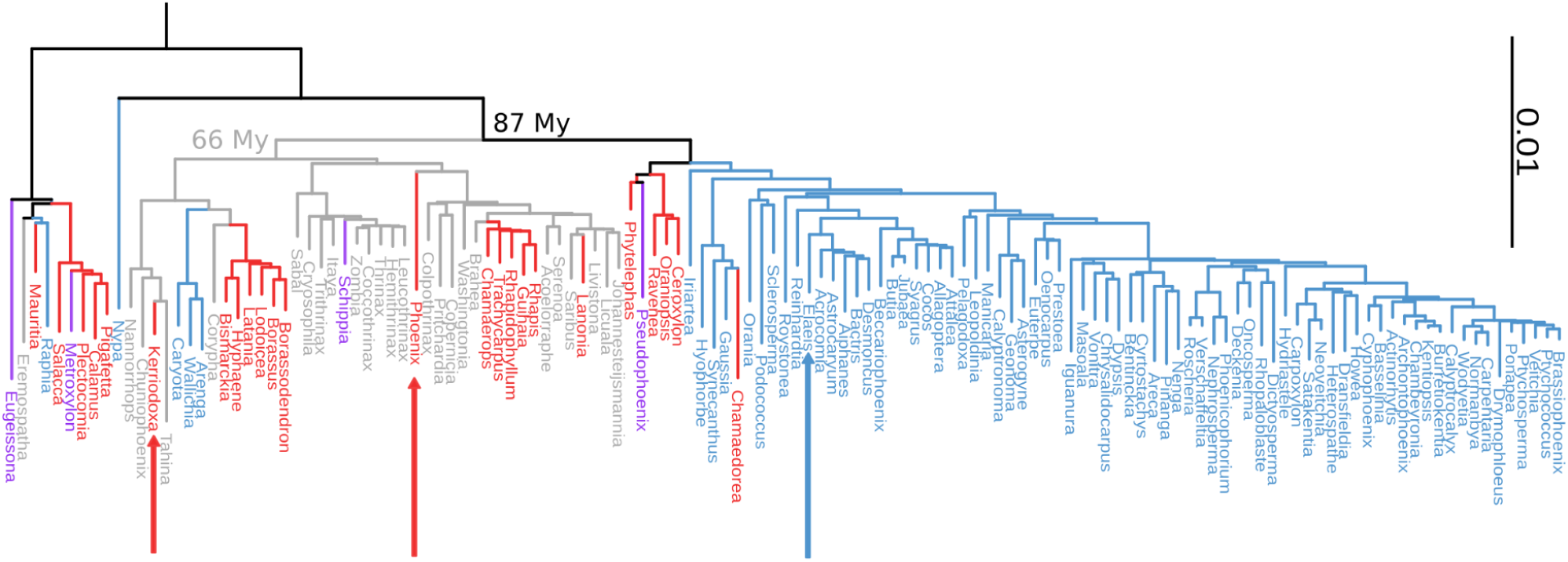
Representation of the sexual system diversity within the palm family in a phylogenetic framework. The plastid phylogeny of palms from Yao *et al*. (2023) is colored according to the sexual system, as classified by Nadot *et al*. (2016) and Cássia-Silva *et al*. (2021): monoecy (blue), dioecy (red), hermaphroditism (grey), and polygamy (purple). Black branches indicate inconclusive ancestral state reconstruction. The branch length corresponds to the number of the estimated substitutions per base (Yao *et al*., 2023). The estimates of the age of the hermaphroditic most recent common ancestor (MRCA) of *Phoenix dactylifera* and *Kerriodoxa elegans* as well as their MRCA with *Elaeis guineensis* (unresolved sexual system, either hermaphroditic or monoecious) are based on Baker & Couvreur (2011). The *Phoenix* and *Kerriodoxa* genera are indicated by red arrows and *Elaeis* genus by a blue arrow.

Sex chromosomes are known to occur in the date palm *Phoenix dactylifera* (Siljak-Yakovlev *et al*., 1996; Cherif *et al*., 2013; Mathew *et al*., 2014), and are shared across its genus (Cherif *et al*., 2016; Torres *et al*., 2018). Despite their relatively old age compared to other dioecious species with sex chromosomes (Renner & Müller, 2021), their non-recombining region has remained small, and the chromosomes have not evolved into a distinctly heteromorphic pair. Torres *et al*. (2018) identified two genes involved in male organ development absent from the X chromosomes, along with another gene that could inhibit female organ development. They suggested that this scenario fits the “two linked sterility mutations” model (Charlesworth & Charlesworth, 1978), with dioecy evolving through a gynodioecious intermediate. However, gynodioecy is absent or very rare in palms (Nadot *et al*., 2016), and the function of the identified genes has not been demonstrated in date palm. Evidence for genetic sex determination has also been found in *Trachycarpus fortunei*, where its XY chromosome system seems to be quite young and unstable (Jousson *et al*., 2023), and possibly in *Lodoicea maldivica* (Morgan *et al*., 2020).

Here we focus on the dioecious species *Kerriodoxa elegans*, for which sex chromosomes have not been previously described, and compare its sex determination with that of the date palm. *K. elegans* is the only species of its genus and was discovered rather recently in Phuket island in Thailand (Dransfield, 1983). According to ancestral state reconstructions, *K. elegans* evolved dioecy independently from the genus *Phoenix*, most probably from a hermaphroditic ancestor (Figure **1**; Nadot *et al*., 2016; Cássia-Silva *et al*., 2021). For comparative analyses on the scale of the palm family, we developed two sequence capture kits targeting exons genome-wide as well as known sex-linked genes, applicable across the palm family. As coding sequences are relatively well conserved among distantly related species (Heyduk *et al*., 2016; Comer *et al*., 2016), this approach allows the direct comparison of sex-linked genes, their evolution and hypothetical function (cf Prentout *et al*., 2021). Sex-linkage was identified through the probabilistic approach SDpop, which is based on the detection of genotype frequencies typical of recombination suppression (Käfer *et al*., 2021) and has been successfully applied to long-lived species that cannot be easily bred in the lab (e.g. *Amborella*, Käfer *et al*., 2022). This approach also allows reconstructing the consensus sequences of the X and Y copies of the sex-linked genes, which were subsequently used for phylogenetic analysis of these genes. We tested these tools on the known sex chromosomes of *P. dactylifera* before applying them to *K. elegans*. We then showed that *K. elegans* has a XY sex determination system that evolved independently from the one in *P. dactylifera*, but has striking similarities that raise intriguing questions about the evolution of sexual systems in palms.

## Material & Methods

### Plant material and DNA extraction

Leaf samples of 10 females and 10 males of *P. dactylifera* were collected in Al Ain, United Arab Emirates and 14 females and 11 males of *K. elegans* in Puerto Rico (Central Cordillera) and in Thailand (Nong Nooch Tropical Botanical Garden). DNA was extracted from 15 mg of dried leaves with the Chemagic DNA Plant Kit (Perkin Elmer Chemagen, Baesweller, DE, Part # CMG-194), according to the manufacturer’s instructions. The protocol was adapted to the use of the KingFisher Flex™ (Thermo Fisher Scientific, Waltham, MA, USA) automated DNA purification workstation.

### Targets and baits design

We designed two separate sets of capture targets. The first one, SexPhoenix (SP), was developed to focus primarily on sex-determining regions. This set includes potential sex-linked genes of *P. dactylifera* (Al-Dous *et al*., 2011; Cherif *et al*., 2016; Torres *et al*., 2018). To enable comparative analysis with these genes, we added nuclear genes unrelated to sex (e.g. salt tolerance genes), chloroplast genes (e.g. *pbsZ* an*d trnG),* and highly conserved low-copy protein-coding nuclear and plastid genes from Heyduk *et al*. (2016). In total, SexPhoenix target set comprises 920 regions (File **S1**). To identify additional potentially sex-linked genes not previously characterized, we developed PalmExon (PE) set, using the transcriptomic data of *P. dactylifera* (NCBI Bioproject PRJNA83433). This target set captures two exonic regions of 30,000 protein-coding nuclear genes (File **S2)**.

From the SexPhoenix and PalmExon target sets, 38,884 and 60,000 enrichment baits were designed, respectively, with a 30-base overlap between baits in SexPhoenix. These consist of biotinylated RNA probes of 90-mers (SexPhoenix) and 80-mers (PalmExon) synthesized by MYcroarray (Arbor Biosciences, MI, USA).

### Target enrichment

For each sample, 100 ng of total DNA were shredded using the NEBNext Ultra II FS DNA Modul (New England Biolabs, Ipswich, MA, USA). The fragmented DNA was ligated to the adapters and the genomic library was constructed following published protocols (Rohland & Reich, 2012; Mascher *et al*., 2013) with some modifications. Two hybridization reactions were implemented, one for each bait kit. In each hybridization reaction, a maximum of 24 samples were multiplexed into one pool, giving a total of 2 pools per reaction. All hybridization steps followed the myBaits protocol version 4.01 (http://www.mycroarray.com).

The final pooled library was sequenced using the Illumina paired-end protocol (2×150bp) on HiSeq3000 sequencer (Novogene, Beijing, China). The detailed protocol for sequence capture is presented in the Supplementary notes.

### Mapping and genotyping

For comparison with the capture kits, we downloaded whole genome sequencing (WGS) data of 20 female accessions and 12 male accessions of *P. dactylifera* from NCBI (bioproject PRJNA505141, Hazzouri *et al*., 2019). In total we analyze five different datasets: (i) WGS *P. dactylifera*, (ii) PalmExons (PE) *P. dactylifera*, (iii) SexPhoenix (SP) *P. dactylifera*, (iv) PalmExons (PE) *K. elegans* and (v) SexPhoenix (SP) *K. elegans*. We used 4 genome assemblies for our analysis, 2 of which from *P. dactylifera* : the homogametic (XX) female DPV01 (Al-Mssallem *et al*., 2013) and heterogametic (XY) male PDMBC4 (Hazzouri *et al*., 2019). The last two are from *Elaeis guineensis* (oil palm), the EG5 (Singh *et al*., 2013) and EGPMv6 (Ong *et al*., 2020) genomes. EGPMv6 is the only one not annotated.

A visual summary of the detection and study of sex-linked sequences is presented in Figure **S1**. For the detection of XY polymorphisms, we used the female (XX) DPV01 genome of *P. dactylifera* as a reference for mapping. In addition to the fact that XY polymorphisms can be detected more reliably than X- or Y-hemizygosity (Käfer *et al*., 2021), assembled genomes from the heterogametic sex might be partially chimeric in the non-recombining region due to the known difficulties of assembling them (Tomaszkiewicz M *et al*., 2017; cf Käfer *et al*., 2021), making the inference of the gametologous haplotypes more difficult. We used the BWA mem2 algorithm (version 2.2.1; Li & Durbin, 2009; Vasimuddin *et al*., 2019) to map the reads, lowering the penalty for mismatches from 4 to 2 (option -B 2), to allow more divergent reads (Y chromosome, *K. elegans*) to be correctly mapped. For comparison with the most recent chromosome-level assembly (PDMBC4) we identified the position of DPV01 annotation with Liftoff (Shumate & Salzberg, 2021). *E. guineensis* assemblies were used to compare synteny and gene content between genomic regions. The unannotated EGPMv6, was used for synteny analyses, while the EG5 annotation was used to identify the homologous genes between *P. dactylifera* and *E. guineensis*. Liftoff was used to position the *E. guineensis* EG5 and *P. dactylifera* DPV01 annotated genes on the *E. guineensis* EGPMv6 assembly (Ong *et al*., 2020). Y-specific genes not present on the DPV01 assembly were studied by recovering unmapped reads and aligning them on the PDMBC4 assembly.

Coverage depth for each base was calculated using bedtools 2.30 genomecov (Quinlan & Hall, 2010). We calculated the mean depth for 100 base windows for all samples, as well as the average for each sex. To calculate the mean depth per gene, we used the windows with at least 50% overlap with genes CDS. To calculate the proportion of female reads for each 100 base windows, we accounted for possible read depth differences between samples by normalizing the depth of each individual. To do so, for each window, the mean depth was divided by the median of the mean depth of all windows of each individual.

SNP calling was done with bcftools mpileup (version 1.9; Danecek *et al*., 2011; Lefouili & Nam, 2022). Genotypes with fewer than 7 reads for the capture data and 10 for the WGS data were discarded . The resulting vcf file was then filtered to keep only the sites within coding sequences according to the DPV01 annotation. We conducted a Principal Component Analysis (PCA) of the genetic variation between individuals using plink 1.90 (Purcell *et al*., 2007) to check population structure and ensure that individuals were not clustered by sex, which could bias the analysis. Additionally, we estimated relatedness between the individuals using the option “relatedness2” in vcftools (Manichaikul *et al*., 2010): values close to 0.5 indicate clones, around 0.25 full siblings, and around 0.125 2nd-order relatives. Paralogous sequences can lead to relatedness overestimation as they harbor heterozygous sites across individuals. As SDpop detects potentially paralogous sequences (see below), if a dataset for a species had high paralogy rate we removed genes with paralogous posterior probabilities above 0.8 and re-estimated the relatedness.

### Sex-linked genes

We used SDpop (Käfer *et al*., 2021) to infer the presence and type (XY or ZW) of sex chromosomes, as well as to identify the sex-linked genes. The tool formalizes the population genetic concepts behind commonly used methods for sex-linkage detection such as male vs female *F_ST_* or GWAS, and puts them in a probabilistic framework that accounts for some frequently encountered sources of error (genotyping errors, paralogy and haploidy). It allows to detect sex linkage (i.e. gametology, XY or ZW, or hemizygosity, when Y or W copies are absent), by inferring the allele frequencies on the putative X and Y copies of the gene from the observed genotype frequencies among females and males. Each gene is assigned a posterior score for each of the segregation types (autosomal, paralog, haploid, XY and X-hemizygous). Genes with a posterior score for XY higher than 0.8 were considered putatively sex-linked. The estimated allele frequencies on the X and Y chromosomes are used by SDpop to reconstruct the X and Y consensus haplotypes of the gametologous genes. For this, we used the SDpop default parameters: if the estimated allele frequency was above 0.95, the allele was considered fixed; if it ranged between 0.6 and 0.95, the major allele was output; and sites with no allele at a frequency above 0.6 were considered missing (“N”).

As SDpop’s posterior scores for a gene are only based on the average allele-genotype frequency equilibria and not on SNP density, we used the latter to refine and confirm SDpop’s results. First, we calculated the average per-gene heterozygosity of the individuals of each sex. As a statistic, we used the proportion of average male heterozygosity per gene, i.e. the average male heterozygosity divided by the sum of average female and average male heterozygosity. This proportion was plotted per gene, except for genes with no heterozygous positions.

Second, codeML (PAML version 1.9; Yang, 2007) was used to infer pairwise synonymous divergence (dS) between the inferred X and Y haplotypes. Sex-specific deviations in genotype frequencies can arise by chance or through linkage disequilibrium with the non-recombining region, but such genes will not have accumulated substitutions independently. Thus, we compared the dS of the putative sex-linked genes to the genome-wide heterozygosity in each species (using the median value of per-gene heterozygosity, excluding genes with paralogous or XY posterior scores over 0.8).

For comparative purposes, we focused on all genes that were putatively sex-linked (i.e. SDpop posterior XY score > 0.8) in at least one dataset, and categorized them as follows. Among the genes with posterior XY scores over 0.8, those that had a dS higher than the species’ median heterozygosity were considered sex-linked with “high confidence” (HCSL), others were considered “probably sex-linked” (PSL). Equally, genes that had posterior XY scores between 0.5 and 0.8 were considered “probably sex-linked”. Genes with autosomal posterior scores above 0.8 were considered as “not sex-linked”. Finally, if genes had on average fewer than 10 individuals genotyped per site, we could not assert sex-linkage or its absence and considered them “inconclusive”. All genes not fitting any of the previous criteria were classified in the inconclusive category as well. These criteria are summarised in Table **S1**. When a gene was present in multiple datasets of the same species, we based this classification either on the dataset with most polymorphisms or, if equal, with more individuals genotyped on average.

We plotted the HCSL genes on the genomes of *P. dactylifera* (PDMBC4) and *E. guineensis* (EGPMv6), which are the most contiguous genomes assembled to date. Our aim was to determine whether these genes are clustered in a single genomic region or dispersed and to identify their syntenic region(s), if any, in a monoecious species (*E. guineensis*). To achieve this, we extracted coding sequences (CDS) from the reference genomes using gffread v.0.12.7 (Pertea & Pertea, 2020). The genome annotation files were converted from GFF to BED format and processed using JCVI functions in Python v. 3.12.7. Pairwise synteny analysis and visualization were performed using MCScan (Python version) (https://github.com/tanghaibao/jcvi/wiki/MCScan-(Python-version)).

### Gene divergence and phylogeny

We identified genes in the *E. guineensis* genome that are homologous to inferred sex-linked genes by selecting the best hit in a blast search against NCBI’s core nucleotide database. Sequences were aligned using muscle (Edgar, 2004), or PRANK (Löytynoja A, 2014) for alignments containing indels shifting the reading frame. Despite the absence of homologous recombination during meiosis, ectopic gene conversion could prevent divergence of the two gametologous copies (Jeffreys & May, 2004; Rosser *et al*., 2009; Niederstätter *et al*., 2013). Geneconv (Sawyer, 1989) was therefore used to detect gene conversion in these alignments. Geneconv identifies stretches with reduced divergence in alignments, indicative of recombination through gene conversion. Geneconv then calculates two p-values, simulated p-values are considered more accurate yet less conservative than Karlin-Altschul (KA) p-values. In order to avoid underestimating the age of recombination arrest of genes, we considered pairwise or global inner-fragments (respectively from one-to-one or one-to-all comparison among the sequences in the alignments) to have undergone gene conversion when at least one of the two p-values was lower than 0.05 between X and Y sequences from the same dataset. For *P. dactylifera*, the same fragment (same start and end position) of the gene was required to fit these criteria for both the Whole-

### Genome Sequencing (WGS) and targeted-capture datasets

The phylogeny was then inferred using BEAST2 v2.7.5 (Bouckaert *et al*., 2019) with a GTR + Ɣ model with 4 gamma categories (Lanave *et al*., 1984; Yang, 1993, 1994a, 1994b). The length of the chain was 250,000,000 steps and a sample was taken every 100 steps after a pre burn-in of 250 samples. A relaxed clock model was chosen (Drummond *et al*., 2006). The prior for the most recent common ancestor (MRCA) age for the root was set to normal distribution with a mean of 87.025 and a standard deviation of 0.625 fitting the 95% confidence interval of this node age in previous dated phylogenies (Baker & Couvreur, 2013). Other parameters and priors were not modified. If convergence of the model was not reached, the length of the chain was increased and the program stopped after all parameters converged and reached an effective sample size (ESS) of 400 on Tracer v1.7.2 (Rambaut *et al*., 2018). A maximum clade credibility tree was then generated with 10% burn in with node heights corresponding to MRCA age (Bouckaert *et al*., 2019). The figures of the inferred trees were generated using ggtree v3.11.1 (Yu, 2020). We additionally used Yao *et al*. (2023) ‘complete-105 regions matrix.300_105_result’ plastid phylogenetic tree to recreate a genus-level phylogeny of palm sexual systems (Nadot *et al*., 2016; Cássia-Silva *et al*., 2021) for context.

### Gene Ontology enrichment

We first updated the Gene Ontology (GO) annotation of the *P. dactylifera* genome (DPV01). Protein sequences were extracted from the genome using GffRead (Pertea & Pertea, 2020) and a functional annotation was elaborated using Fantasia with default parameters (Martínez-Redondo *et al*., 2024). GO terms were filtered and manually curated based on scores calculated by Fantasia (filtration threshold of 0.3).

To investigate the enrichment of sex-related GO terms among genes identified in the suggested sex-linked region (see Results), we conducted a GO enrichment analysis using the topGO package in R version 2.58 (Alexa & Rahnenfuhrer, 2010). We considered the genes of interest, the set of 92 genes located in the proposed sex-linked region. Two distinct gene lists were used as reference universes: (i) all genes captured by the two sets of baits, which had at least one read coverage after mapping sequencing data to the targets, and (ii) genes with a minimum of seven reads coverage on the targets. The GO enrichment was elaborated for the go-category, biological process, using the topGO weight01 algorithm and the Fisher statistical test. The P-value < 0.05 was set as the cutoff for the significance of the gene enrichment. We note that the weight01 algorithm is an improved method that combines the elim and weight algorithms (Alexa *et al*., 2006; Grossmann *et al*., 2007).

## Results

### Characterizing sex-linked genes in *P. dactylifera*

In *P. dactylifera,* the average mapping rates for the whole-genome sequencing data (WGS, Table **S2**), and SexPhoenix (SP) and PalmExons (PE) capture kits (Table **S3**), were 94.28%, 99.46% and 99.51%, respectively. Target capture covered 779 of the 832 targeted regions at a depth of over 1X with a median depth of 38.84 reads per base and a median coverage of 38.57% of the targets for SP. The PE kit captured 25,579 of the 29,983 targeted regions at a depth of over 1X with a median depth of 6.49 reads per base and a median coverage of 11.35% of the targets. Due to high relatedness between samples, we discarded three female and three male samples from the WGS data (Figure **S2a**,**S3a**, Table **S2**), and one male sample for the sequence capture data (Fig **S2b**,**S3b**, Table **S3**). Therefore, for the WGS data, the datasets consisted of 17 females and 9 males, while for both capture data (SP and PE), 9 individuals of each sex were included in the datasets analyses. After filtering by read depth (over 10 reads per site per individual for WGS data, over 7 for the capture kits), we analyzed over 450,000 SNPs and indels for the sex-linkage analysis in 25,100 genes (98% of the annotated protein-coding genes in the female date palm genome, DPV01, median gene length coverage over 99%) using the WGS data (Table **S4**). For the capture data, a total of 4,693 variant positions in the SP data covering 412 genes (1.6% of the annotated genes, median gene length coverage 75%) and 180,035 SNPs in the PE data covering 19,889 genes (77%, median gene length coverage 48%) were analyzed (Table **S4**).

Upon analysis of the WGS data with SDpop, 90% of the polymorphic sites were inferred to be autosomal (Table **S5**) while the proportions of haploid and sex-linked (XY gametologous and X-hemizygote) sites were inferred to be fewer than 1%. In the sequence capture datasets, fewer than 1% of sites were found sex-linked in PalmExons (Table **S5**), consistent with the WGS data proportions. As expected by design of the capture kits, the proportion of sex-linked sites within SexPhoenix baits was much higher, 12% (Table **S5**). Excluding genes with paralogous or XY posterior scores above 0.8 in SDpop, median per gene heterozygosity was 0.0021 for both WGS data (24,472 genes) and PE data (19,703 genes). We hence used 0.0021 as the synonymous divergence (dS) threshold value for high-confidence sex-linked genes (HCSL) in *P. dactylifera* (Table **S1**).

The highest XY scores inferred by SDpop for the *P. dactylifera* genes on the PDMBC4 genome (Fig. **2a**, outer ring) are on chromosome 12, the previously identified sex chromosome (Mathew *et al*., 2014) (Fig. **2b**). A 3 Mb region between 9.4 Mb and 12.4 Mb on this chromosome mostly contains genes with high XY scores, while containing no genes for which autosomal segregation was inferred (Fig. **2d**). The location of the non-recombining region is also visible in a comparison of average female and male heterozygosity, with a strong increase in male heterozygosity as expected in an XY system (Fig. **S4**). We combined the results of WGS, PE and SP datasets by using the one with the most polymorphisms when a gene was present in multiple datasets (see Material and Methods). We thus classified 45 genes as high-confidence sex-linked (HCSL), because they had SDpop XY posterior scores over 0.8 and X-Y dS higher than the average nucleotide diversity (0.0021, Table **S1**), both criteria being strong indicators of recombination suppression. We additionally classified 42 genes as probably sex-linked (PSL) when their dS was lower or when their probability to be XY was between 0.5 and 0.8. Among HCSL genes, 13 are on chromosome 12 and 26 on PDMBC4 unassigned scaffolds (File **S3**). Similarly, 15 PSL genes are on chromosome 12 and 18 on unplaced scaffolds (File **S3**). Few genes had posterior scores higher than 0.8 for X-hemizygosity (WGS, PE and SP), but upon verification, none of these genes had significant differences in female and male read coverage (File **S4**), so we conclude no X-hemizygous genes were detected.

**Figure 2.**
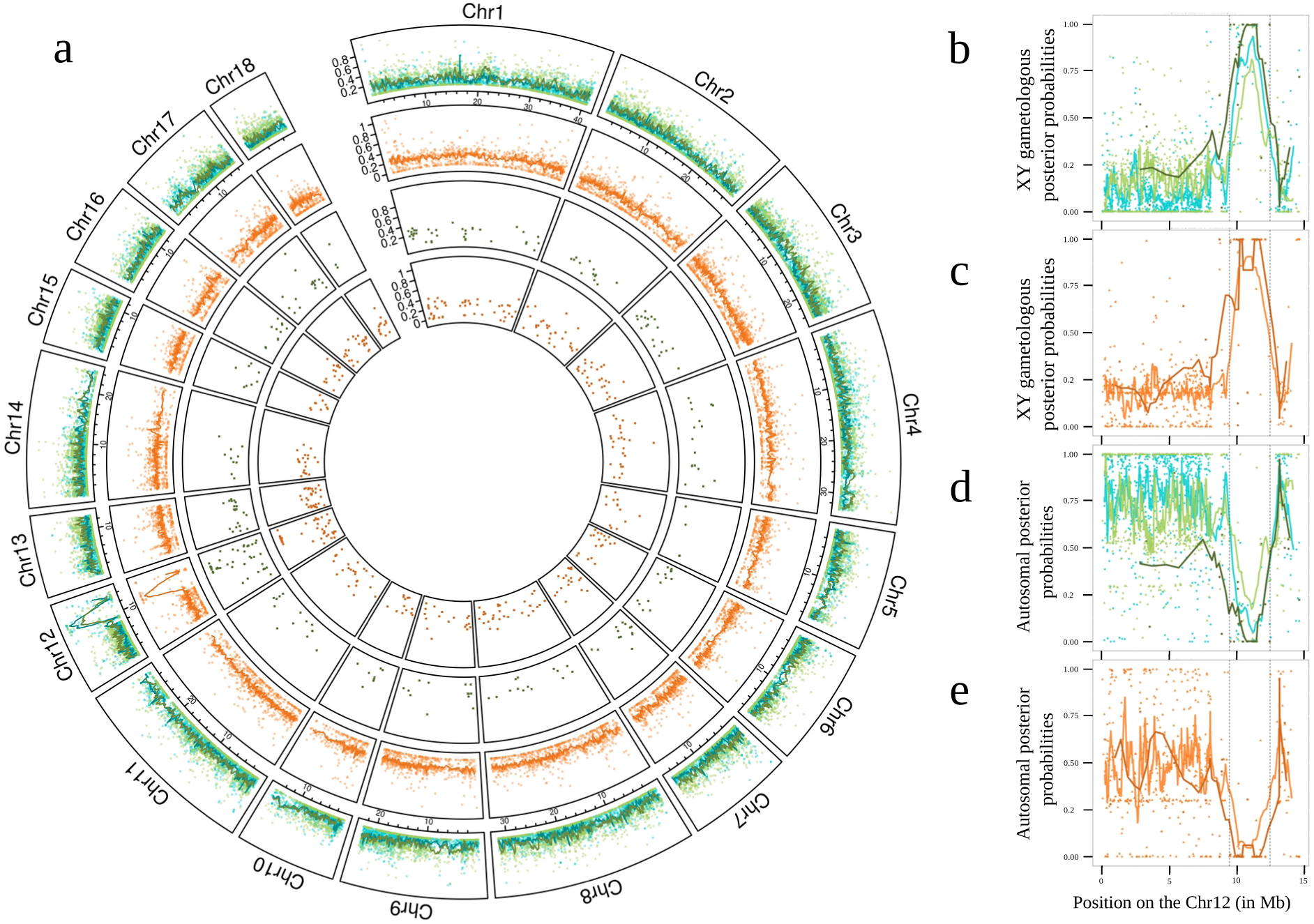
Probability of sex linkage (XY) of each gene for *Phoenix dactylifera* and *Kerriodoxa elegans*. The position of the genes is taken from the PDMBC4 assembly of *P. dactylifera* (Hazzouri *et al*., 2019). *P. dactylifera* is indicated in blue (whole genome resequencing data) and green (capture kit data), *K. elegans* in orange. (**a**) Genome-wide distribution of XY posterior probabilities obtained with SDpop. Outer two rings show whole-genome resequencing and capture (PalmExons) data, the inner two the SexPhoenix capture kit. **(b,c)** XY posterior probabilities per gene on chromosome 12. **(d,e)** Autosomal posterior probabilities per gene on chromosome 12. Lines show the rolling mean with a window size of fifteen genes for the PalmExons kit (light green or orange) and whole genome resequencing data (blue) and five genes for the SexPhoenix kit (dark green or orange).

Among the genes previously described as Y-specific in *P. dactylifera* (Torres *et al*., 2018), only LOG3like (LOC103701078) was inferred as XY in our analysis (File **S3**). Further investigation confirmed this was due to paralogy of the Y-specific gene and an autosomal gene, as previously found (Torres *et al*., 2018): males had more reads than females and reconstructed X and Y haplotypes aligned to two different loci in the date palm genome (LOC120104231 on chromosome 17 in the PDMBC4 assembly for the X sequence, and mRNA from the Y chromosome (MH685630) for the Y sequence). The duplication between the LOG autosomal and Y-linked copies is estimated at 53.76 My (Figure **S5**), likely predating sex-linkage. Concerning the other two Y-specific genes identified previously, we confirmed by remapping previously unmapped reads (i.e. those that did not align to the female DPV01 genome) to the PDMBC4 genome that both GPAT3 (LOC120108105) and CYP703 (LOC120107104) had no female reads for *P. dactylifera* (File **S3**). Thus, our approach based on target capture sequencing and sex-linkage identification in *P. dactylifera* using SDpop leads to similar results as previous studies.

### *K. elegans* and *P. dactylifera* sex chromosomes share sex-linked genes and a common genomic region

The capture sequences of *K. elegans* had 83.95% of properly-paired reads for SP and 99.20% for PE when mapped on the DPV01 female *P. dactylifera* genome. Most *K. elegans* samples were related, with initial analyses suggesting they are full siblings (Figure **S2b,S3b**), but after removing possibly paralogous genes from the data (genes with SDpop’s paralogous posterior scores over 0.8), most individuals were inferred to be first-degree cousins (Figure **S6**). Some inbreeding is expected as *K. elegans* has small populations and many samples come from botanical gardens. This will have a negligible effect on the inference of sex-linkage as relatedness was found between female and male individuals alike, and our inferences were based on population genetic equilibria as well as dS between X- and Y-copies, and the latter is not affected by relatedness. SDpop was run on 14 females and 11 males of *K. elegans* with 161,384 variant positions for PE covering 15,051 genes (58% of the DPV01 annotated genes, median gene length coverage 43%) and 8,395 variant positions covering 655 genes (2.5%, median gene length coverage 54%) for SP (Tables **S3**,**S4**).

As sex chromosomes were not previously known in *K. elegans*, we ran SDpop’s models for XY sex chromosomes, ZW chromosomes and without sex-linkage. The model that most accurately fitted both datasets of *K. elegans* according to the Bayesian Information Criterion (BIC) is XY (Table **S5**), i.e. the same as *P. dactylifera*. Median heterozygosity per gene was 0.0011 in PE data excluding genes with XY or paralogous scores above 0.8 (13,442 genes). Thus, 0.0011 was used as the dS threshold value for *K. elegans* HCSL genes (Table **S1**).

When a gene was present in both datasets, we used the data from the dataset in which it had the most polymorphisms for subsequent analysis. We identified 59 HCSL genes in *K. elegans*, 25 on chromosome 12, 4 on other linkage groups, 29 on unplaced scaffolds and 1 not positioned on the PDMBC4 assembly (Figure **2**, File **S3**). SD-pop analysis suggested that some genes could be X-hemizygous but after analysis of the female and male coverage, this could not be confirmed (File **S4**). Most genes found to be sex-linked in *P. dactylifera* and *K. elegans* were common between the two species, even those on unplaced scaffolds (Figure **2**, Figure **S4**, File **S3**). More precisely, of the 45 genes inferred as HCSL in *P. dactylifera* and the 59 in *K. elegans*, 21 were so in both species, whereas only 2 HCSL genes in *P. dactylifera* and 10 in *K. elegans* were found to be autosomal in the other species (File **S3**). Furthermore, 15 HCSL genes in *K. elegans* were probably sex-linked (PSL) in *P. dactylifera*, and three HCSL genes in *P. dactylifera* were PSL in *K. elegans* (Fig. **3**, File **S3**). Mapping previously unmapped reads to the PDMBC4 male genome revealed that two Y-specific genes of *P. dactylifera*, GPAT3 (LOC120108105) and CYP703 (LOC120107104), were XY in *K. elegans*. The third Y-specific gene of *P. dactylifera*, LOG3like (LOC103701078), was inferred as autosomal in *K. elegans* (File **S3**).

**Figure 3.**
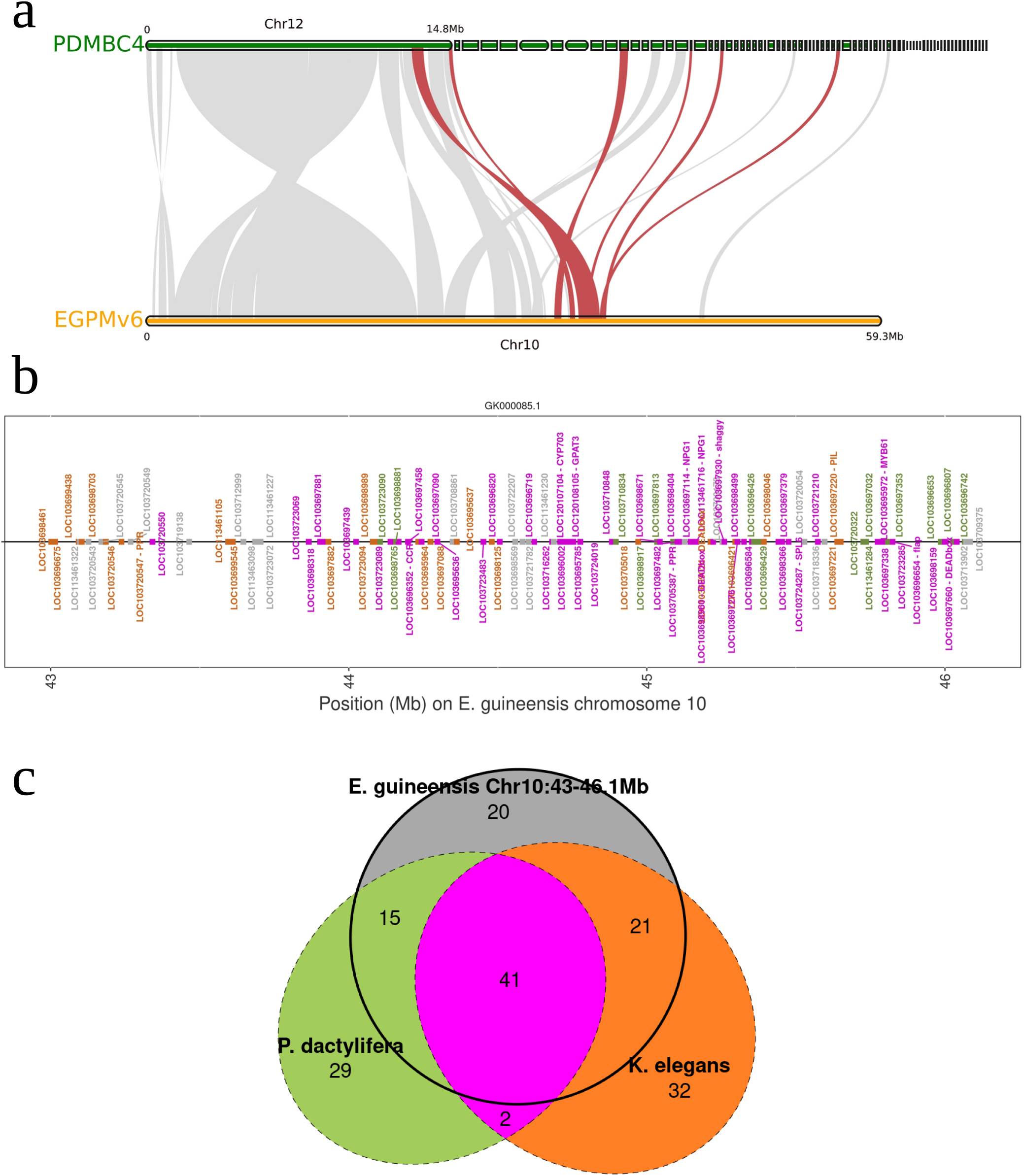
Position of *Kerriodoxa elegans* and *Phoenix dactylifera* sex-linked genes on *Elaeis guineensis* chromosome 10. Genes with posterior probability to be XY above 0.8 in at least one dataset or present between 43 and 46.1Mb on chromosome 10 of the *E. guineensis* EGPMv6 genome were categorized as follows: high-confidence sex-linked, probably sex-linked, inconclusive, not sex-linked or missing data. **(a)** Synteny between *E. guineensis* EGPMv6 chromosome 10 (orange) and *P. dactylifera* PDMBC4 chromosome 12 and several unplaced scaffolds (green). Synteny of sex-linked genes is colored in red. **(b)** Positions of genes between 43 and 46.1Mb on chromosome 10 of *E. guineensis*. Sex-linked genes are colored in green for *P. dactylifera*, orange for *K. elegans*, and magenta for both. Genes from other categories (i.e. either inconclusive, or absent in at least one species, or autosomal in at most one) are colored in grey. **(c)** Euler diagram of the number of sex-linked genes (HCSL and PSL) in *P. dactylifera* (green) or *K. elegans* (orange) or positioned between 43 and 46.1Mb on the chromosome 10 of *E. guineensis* (grey).

As the PDMBC4 assembly was generated using a male individual, a considerable fraction of the non-recombining region might have been difficult to assemble. However, the sex-linked region in *P. dactylifera* shares synteny with a genomic region on the monoecious oil palm ( *Elaeis guineensis*) chromosome 10 (Fig. 3a), so we positioned the sex-linked genes of *P. dactylifera* and *K. elegans* on the *E. guineensis* EGPMv6 genome. They indeed fall in a single region between 43 and 46.1 Mb of the *E. guineensis* chromosome 10, in a more contiguous pattern than in PDMBC4 genome (Fig. **3**, table **S6**). In total, this region contains 108 protein-coding genes according to the EG5 annotation for *E. guineensis* and 95 according to the DPV01 annotation for *P. dactylifera,* with the addition of two Y-specific genes from the PDMBC4 annotation (Figure **S7**). It contains 78% of genes inferred as HCSL in either *P. dactylifera* or *K. elegans*, including 90% of common HCSL genes. None of the genes in this region were inferred to be autosomal in *K. elegans*, while 17 were in *P. dactylifera* (Fig. **3**, Table **S7**, File **S3**), indicating that the sex-linked region in *K. elegans* might be slightly larger.

To determine if the sex-linked region was enriched in genes related to sex determination, we first improved the functional annotation of the DPV01 genome assembly. The updated annotation resulted in the annotation of 24,625 genes (ca. 80% of the genome). The *de-novo* functional annotation was compared and merged with the annotation of *P. dactylifera* PDMBC4 genome. The final annotation encompasses GO IDs for 29,167 genes, covering approximately 95% of the DPV01 genome (File **S5**). We then performed GO enrichment analysis on all 92 protein-coding genes between 43.3Mb and 46.1Mb on chromosome 10 of *E. guineensis* (excluding the 43Mb to 43.3Mb region as it contains genes that are only sex-linked in *K. elegans*; Fig. **3b**, File **S3**). The analysis showed highly similar results when taking into consideration the genome-wide captured genes (universe genes) with 1X and 7X read coverage (File **S6**). We identified a significant enrichment of GO terms (p < 0.05) covering various biological processes, including DNA repair, stress responses, and protein translation (File **S6**). While a few GO terms related to plant reproductive mechanisms, such as pollen tube adhesion, plant ovule development, specification of floral organ identity, and auxin-activated signaling pathway, showed significant enrichment (p < 0.05), the copy number of genes associated with these terms was not overrepresented which reduces the statistical significance of enrichment for these genes (File **S6**). Therefore, our analysis did not reveal convincing sign of the over-representation of GO terms for genes with floral development or sex-linked functions.

### Independent recombination arrest for *P. dactylifera* and *K. elegans*

To investigate whether recombination suppression took place in a common ancestor of *Kerriodoxa* and *Phoenix*, we inferred the gene trees and divergence times for the 42 genes that were found to be sex-linked in both species (either HCSL in both species or HCSL in one and PSL in the other, see Figure **3b**, File **S4**), using the X and Y consensus haplotypes inferred using SDpop, and *E. guineensis* as an outgroup. Prior to the analysis, we found possible gene conversion in 3 genes in *K. elegans* and 3 different ones in *P. dactylifera* (Table **S8**). The effect of either retaining or removing these stretches on the inferred divergence times is negligible (Figure **S8**); for further analyses, we nevertheless chose to discard these stretches.

In all phylogenetic trees, the X and Y sequences from each dioecious species formed a strongly supported clade (Fig. **4a**, **S5**), indicating that recombination suppression evolved after the divergence of the *Phoenix* and *Kerriodoxa* lineages. The node ages in these 42 sex-linked gene trees, that were calibrated using previously estimated divergence times at the level of the palms family, confirmed that recombination suppression occurred much later than the split between the *Phoenix* and *Kerriodoxa* lineages (Fig. **4b**). The mean age of divergence between *P. dactylifera* and *K. elegans* is 61.86 My, whereas mean ages of X-Y haplotypes splitting for the five most divergent genes of each species were estimated to be 22.07 My for *P. dactylifera* and 19.32 My for *K. elegans* (Fig. **4b**). Thus, despite the similarity in gene content and age of these regions, none of the common genes in the sex-linked region showed any evidence of recombination suppression in the last common ancestor of *P. dactylifera* and *K. elegans*. Among the genes that were sex-linked in only one species, we did not observe substantially higher X-Y dS values than for common sex-linked genes (File **S4**), indicating that none of the species-specific sex-linked genes stopped recombining earlier. We conclude that the sex chromosomes of *P. dactylifera* and *K. elegans* evolved independently, which is congruent with the ancestral state reconstructions of sexual systems (Fig. **1**).

**Figure 4.**
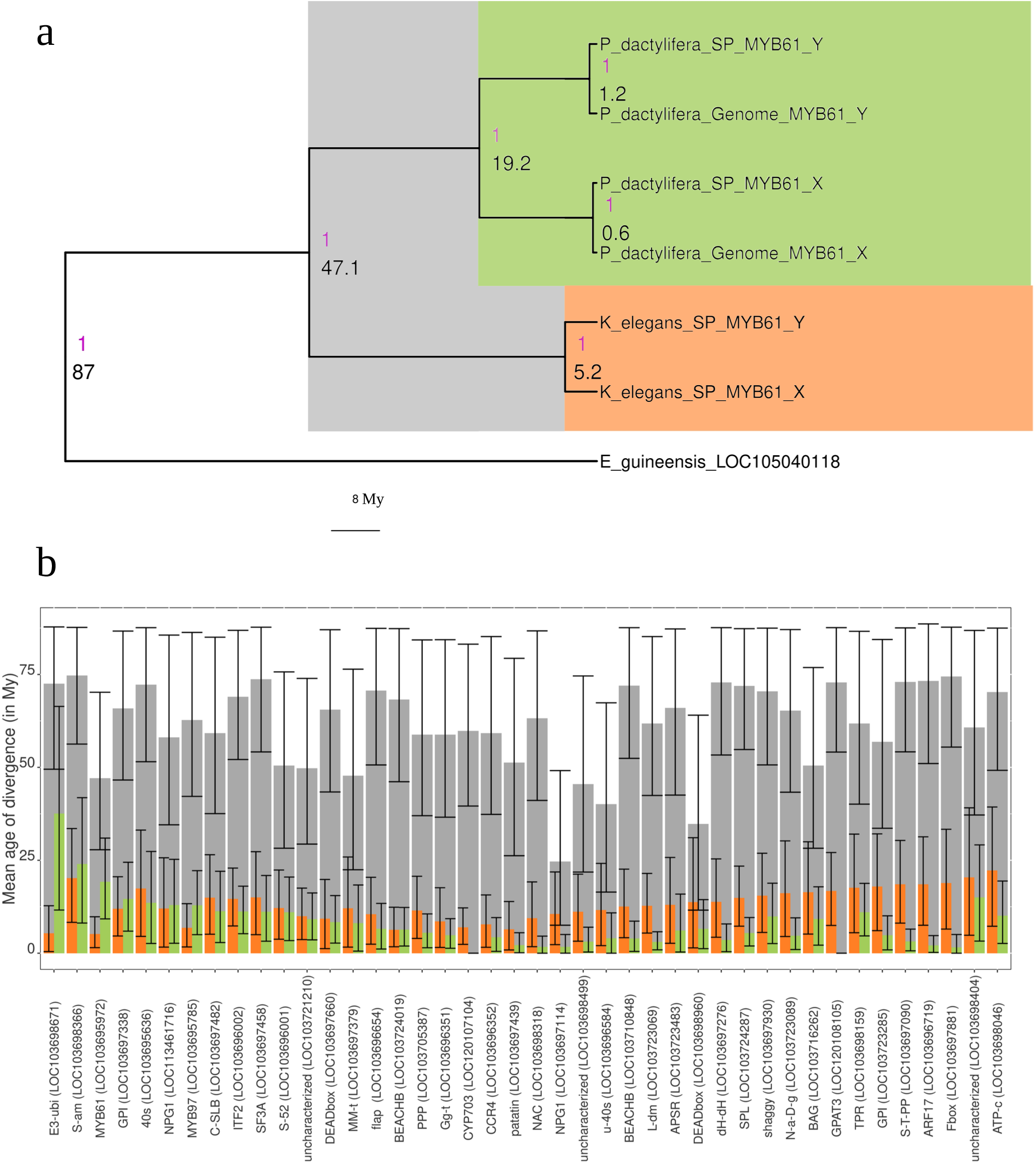
Ages of species and gametolog divergence for common sex-linked genes in *Phoenix dactylifera* and *Kerriodoxa elegans*. Age estimates are colored in grey for the divergence of the *Phoenix* and *Kerriodoxa* lineages, in green for *P. dactylifera* X-Y divergence and in orange for *K. elegans* X-Y divergence. **(a)** Phylogenetic tree for the MYB61 (LOC103695972). Node values correspond to the mean age for the node’s most recent common ancestor (MRCA; black) and the support of the node (magenta). **(b)** Mean age of divergence of X and Y sequences for common sex-linked genes compared to the age of divergence between the MRCA of *P. dactylifera* and *K. elegans*. Error bars correspond to the 95% probability density of the node age.

Synonymous divergence between the X and Y copies of sex-linked genes is rather low, typically lower than 3%, which is strikingly different from a species like *Silene latifolia* which has about 10% synonymous divergence for 11 My of divergence (Krasovec *et al*., 2018). We investigated whether divergence of sex-linked genes with their *E. guineensis* homologs was different than for autosomal genes, using 18 genes that were sex-linked in both species (using SP data which covers a larger part of the genes in this region, Figure **S9**, Table **S4**) as well as ten random autosomal genes from the BUSCO *Liliopsida* database (Table **S9**). Cumulated synonymous divergence (dS) with *E. guineensis* was similar for autosomal and sex-linked genes (Figure **S9**), indicating that the sex-linked genes experienced substitutions at similar rates as autosomal genes. This was expected based on the relatively short time for which these genes have stopped recombining.

## Discussion

### A sex-linked region recruited independently in two distantly related palm species

Based on exon target-capture sequencing and subsequent analyses, we show that *Kerriodoxa elegans* and *Phoenix dactylifera,* two palm species that diverged approximately 66 My ago, have similar XY sex-linked regions. The estimated ages of recombination suppression in both species show that these sex-linked regions evolved independently between 10 and 40 My ago, in line with ancestral state reconstructions of the sexual systems that suggest that both species most likely evolved dioecy independently from hermaphroditism (Figure **1**; Nadot *et al*., 2016; Cássia-Silva *et al*., 2021). Despite their independent origins, *P. dactylifera* and *K. elegans* sex-linked regions exhibit striking similarities, including their size, relatively low divergence considering the age of recombination suppression, and minimal degeneration (no loss of Y genes), although the region might be slightly larger in *K. elegans*. Such convergent evolution seems unique among known plant sex chromosomes.

In animals, gonochorism (sex separation) is ancestral in many taxa, but how sex is determined varies widely, and sex chromosomes have evolved many times independently (Bachtrog *et al*., 2014). Convergent evolution of sex chromosomes has been reported in a few animal clades. For example, in *Xenopus*, it has been suggested that sex chromosome convergence arises due to frequent turnovers, with homologous regions containing key sex-linked genes shared among distant species (Furman & Evans, 2016). Indeed, in animals, gonadal developmental pathways are highly conserved, with similar precursors found in both invertebrates and vertebrates (Kopp, 2012), probably due to the fact that gonochorism is their ancestral state. Consequently, Dmrt and SOX genes have independently evolved multiple times to serve as sex determinants (Kopp, 2012). The mere presence of Dmrt1, or copies of it, on a chromosome makes it likely to evolve into a sex-determining gene, as seen repeatedly in amniotes (O’Meally *et al*., 2012; Ezaz *et al*., 2017).

In plants, dioecy evolved many times independently, and sex determination mechanisms have long been thought to be very diverse (Mitchell & Diggle, 2005; Henry *et al*., 2018). However, as for animals, it is widely accepted that plant sex-determining regions (SDRs) or sex chromosomes contain critical genes involved in sexual differentiation and the development of sex-specific traits. These genes often serve as key regulators in pathways that establish and maintain dioecy, underpinning the evolutionary significance of SDRs in controlling sex-specific phenotypes, and recent progress suggests that several pathways could be common among distantly related plant species (cf Leite Montalvao *et al*., 2021).

### The shared sex-linked region contains genes related to flower sexual development

It seems highly unlikely that a single, 3 Mb region evolved recombination suppression twice just by chance. Recombination suppression is believed to originate with at least one sex-determining gene, implying that the sex-determining genes in both species are located within this region. Which genes, then, are present in this region and could account for its independent recruitment in the two species?

While a GO term analysis of the sex-linked region did not convincingly demonstrate an over-representation of genes involved in plant reproductive development, we identified several potentially interesting genes with functions in flower and sexual organ development based on gene annotations. A substantial number of genes has been found to be involved in pollen and anther development and male sterility in a range of other plants, and a few other genes are involved in ovule development, hormone signaling and inflorescence branching (Table **1**). In *P. dactylifera*, Torres *et al*. (2018) proposed a hypothesis for the evolution of dioecy and sex chromosomes, identifying three Y-specific genes. Among these, *GPAT3*, a glycerol-3-phosphate acyltransferase, and *CYP703*, a member of the cytochrome P450 subfamily, are crucial for anther and pollen development and male fertility in monocot species (Yang X *et al*., 2014; Li & Liu, 2017; Somaratne *et al*., 2017; Xie *et al*., 2018). Another gene, *LOG3*, belongs to the LONELY GUY (LOG) gene family, which is involved in cytokinin activation and ovule development in female flowers (Kurakawa *et al*., 2007; Yamaki *et al*., 2011). Loss of male function in some individuals could have led to gynodioecy (involving recessive loss-of-function of *GPAT3* and/or *CYP703*), followed by the loss of female function (dominant suppression by LOG3) leading to dioecy in date palm (Torres *et al*., 2018). Although our methods were not primarily designed to detect Y-specific genes, additional analyses confirmed these genes are Y-specific in *P. dactylifera*. In *K. elegans*, however, *GPAT3* and *CYP703* were inferred as XY gametologs, and *LOG3* as autosomal. Furthermore, the X-Y copy divergence of these genes in *K. elegans* was not among the highest values found for this species, suggesting that they have been recruited after initial recombination suppression involving the sex-determining gene(s) in this species. While functional analyses of sex-linked genes have yet to be conducted in both species, these findings suggest that the master sex-determining genes in *K. elegans* are different from those in *P. dactylifera*.

**Table 1.**
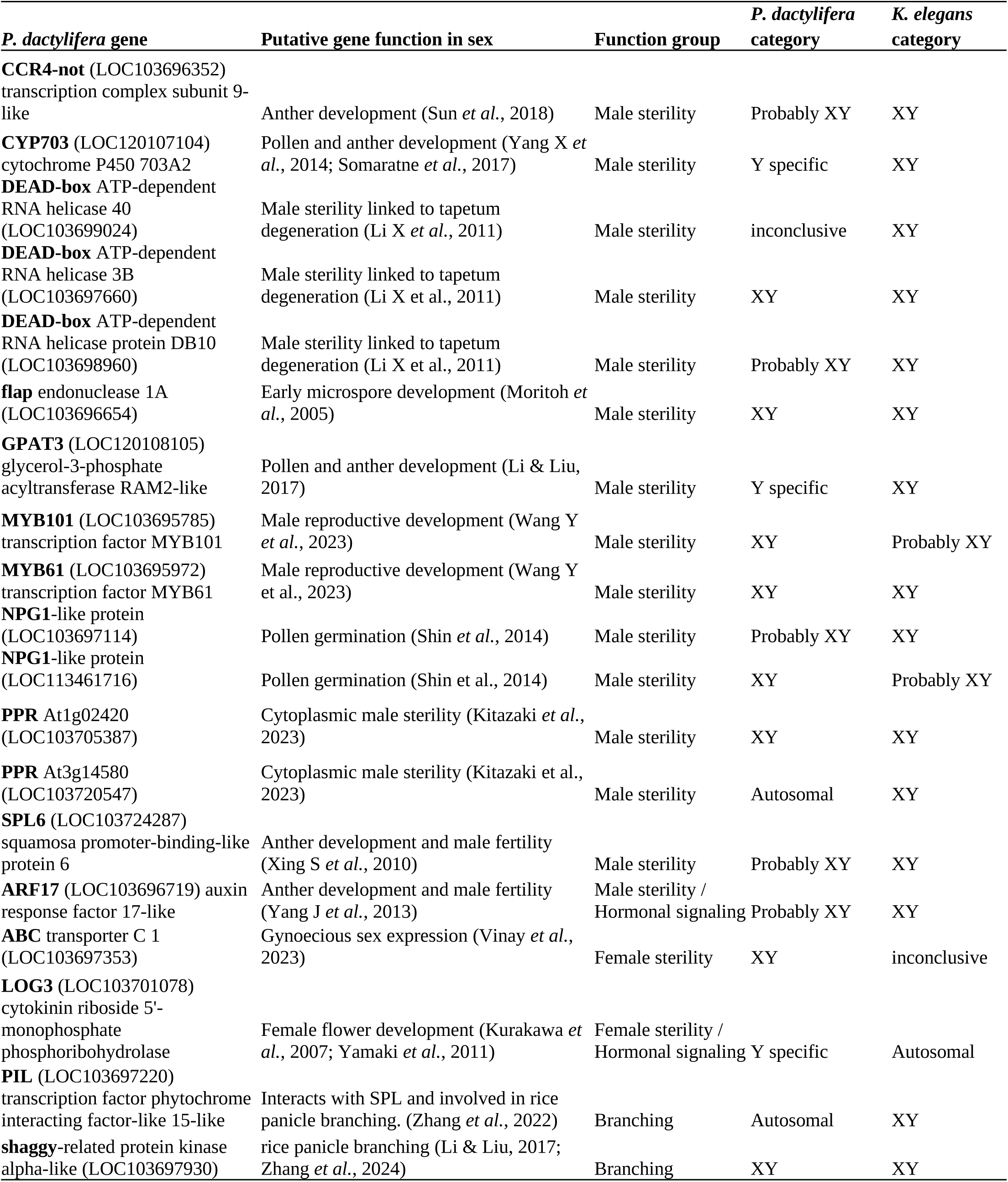
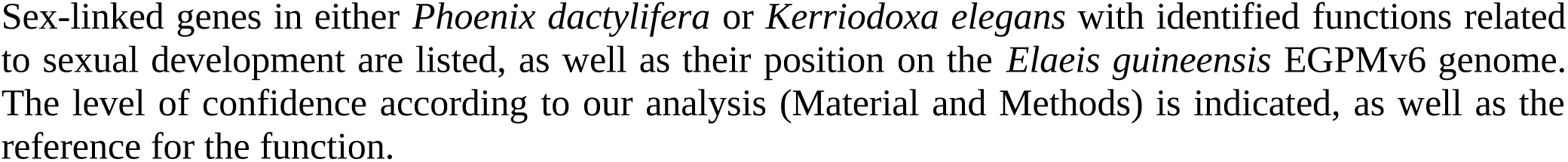
Sex-linked genes with functions related to flower development.

The role of hormones (cytokinins, auxins and jasmonate) in palm sex differentiation has been demonstrated in the monoecious species *Elaeis guineensis* and *Areca catechu* through content variations between female and male inflorescences (Tregear *et al*., 2022; Zhou *et al*., 2022). In *E. guineensis*, an antagonistic regulatory model has been proposed for both sexual differentiation and branching activity in inflorescences, as male inflorescences have more primary branches than female ones (Tregear *et al*., 2022). This model involves the *LOG* gene, the *SQUAMOSA PROMOTER-BINDING PROTEIN-LIKE* (*SPL*) gene, which plays a role in inflorescence development and branching (Kellogg, 2022), and the *PHYTOCHROME-INTERACTING FACTOR-LIKE* (*PIL*) family of transcription factors, which directly interact with SPLs to repress branching and tillering (Zhang *et al*., 2022). *E. guineensis* female inflorescence development is characterized by cytokinin accumulation, increased expression of *LOG* and *SPL* genes, and reduced levels of miRNA regulators, while male inflorescences exhibit higher auxin levels and increased expression of *PIL* genes (Tregear *et al*., 2022). As genes similar to *SPL*, *PIL* and *LOG3* are sex-linked in *P. dactylifera* and/or *K. elegans* (Table **1**), the same genes could be involved in flower sex determination in monoecious palm species and sex determination at the whole-plant level in dioecious species. Not all these genes need to be present in the sex-determining region, as one of them could act as a master-switch gene and also control expression of autosomal genes that are involved in the pathway. The fact that the same genomic region has been recruited twice indicates that it is rich in such potential master-switch genes.

The sex-linked region in *P. dactylifera* and *K. elegans* has conserved synteny since its split with *E. guineensis.* This region has furthermore been shown to be syntenic in two other monoecious species, *Cocos nucifera* and *Areca catechu* (Zhou *et al*., 2022). It was therefore probably present in the common ancestor of the Arecoideae (to which *E. guineensis*, *C. nucifera* and *A. catechu* belong) and the Coryphoideae (with *P. dactylifera* and *K. elegans*) subfamilies, estimated to be 87 My old (Baker & Couvreur, 2013). Considering its composition in sex-related genes, we hypothesize that this region could act as a “genetic toolbox” for the development of unisexual flowers in palms. It could play a key role in facilitating transitions between monoecy, dioecy, and hermaphroditism, even if the master sex-determining genes might be different. It is noteworthy that palms represent one of the plant families with the highest proportion of unisexual flowers, with around 80% of species, compared to only about 10% across angiosperms, and have undergone many transitions between sexual systems (Nadot *et al*., 2016). Genomic colocalization of genes involved in the same pathway could facilitate concerted regulation (Hurst *et al*., 2004) and thus evolutionary transitions involving the pathway. Interestingly, a region responsible for sex determination in *Vitis* was recently shown to constitute a conserved collinear region specific to flowering plants, containing many genes related to flower development (Massonnet *et al*., 2024). Currently, however, and by contrast to the “toolbox” region in palms, it is not known to have given rise to sex chromosomes following other transitions to dioecy.

### Palms as a model system for sex chromosome evolution

Our findings raise the question whether the region we identified in *P. dactylifera* and *K. elegans* might have given rise to sex chromosomes in other palm species. We here also presented the approach that could be used to answer this question. Palms are a large, mostly tropical family for which only a few, economically important species have been studied. They often exhibit large genome sizes: the ancestral genome size was approximately 3.6 Gb, with some species reaching genome sizes as large as 29 Gb (Barrett *et al*., 2019). Despite the increasing accessibility of long-read genome sequencing and assembly, studying many species that cannot be easily bred in the lab remains challenging, especially when genome sizes are large. Sex chromosomes raise particular difficulties associated with divergence, transposable elements, and inversions (Carey *et al*., 2022; Hobza *et al*., 2024). Indeed, we have shown that more than half the genes that are probably part of *P. dactylifera* sex-linked region were not placed on the latest genome pseudomolecules (Hazzouri *et al*., 2019). To investigate sex linkage in multiple dioecious palm species with limited genomic resources, we developed a method based on the reduction of genome complexity (Fig. **S1**). This approach uses sequence capture datasets designed mainly from the female date palm genome (DPV01) and targets either previously identified sex-linked regions (SexPhoenix, SP) or exonic sequences (PalmExon, PE). This latest target capture set provides genomic information on 20,000 genes in *P. dactylifera*, as well as 15,000 genes in *K. elegans*, despite their 66 million years divergence. As coding sequences are generally relatively well conserved at the level of a plant family, this approach allows investigating palm species in a cost-effective manner.

Our approach also incorporates the recently developed tool SDpop, specifically designed to identify and study sex-linkage in non-model species, requiring only a small number of individuals from which material can be collected in the wild, as long as sex can be phenotypically determined (Käfer *et al*., 2021). SDpop assigns a probability to be sex-linked to genes and allows further investigation, by reconstructing their X and Y sequences from which divergence can be inferred. The use of both the posterior scores and X-Y divergence increases the robustness of the method. The genes found to be sex-linked are then positioned onto the date palm genome and genetic map to identify the sex-linked regions and sex chromosomes. The localization of the majority of sex-linked genes within a single genomic region reinforces the confidence in the results. We showed this strategy is effective, successfully reproducing previous findings on *Phoenix dactylifera* sex chromosomes (Mathew *et al*., 2014; Torres *et al*., 2018; Hazzouri *et al*., 2019), and identifying the sex chromosomes in a distantly related species, *Kerriodoxa elegans*, without prior knowledge. This approach does not provide a full picture of the sex chromosomes, as haplotype-aware genome sequencing and assembly would be able to provide, but can be used as a first comparative analysis of dioecious species, giving insights in the number and identity of sex-linked genes and sequence divergence.

Given the importance of several palm species as vital crops, where only females produce fruits and seeds, these genes could be used to develop genetic sex markers that allow early selection of female plants with high added value, thereby avoiding to wait before the sexual phenotype becomes apparent, which can take up to 15 years (Cherif *et al*., 2013). More broadly, the existence of a region determining the unisexuality of inflorescences and the knowledge of the genes it contains offer perspectives for controlling the sex ratio of cultivated palm species, whether they are dioecious or monoecious (Ong *et al*., 2020).

## Acknowledgements

This project (ID 2101-014 and ID 2201-007) was funded through LabEx AGRO 2011-LABX-002 (under I-site Muse framework) coordinated by Agropolis Fondation. We thank Mike Dahme, Central Florida Palm and Cycad Society, USA for providing *Kerriodoxa elegans* material. We also thank the ISO 9001 certified IRD i-Trop HPC (South Green Platform) at IRD Montpellier for providing HPC resources that have contributed to the research results reported within this paper, as well as Audrey Weber from the Genotyping platform, INRAe Arcad AGAP, for sequencing.

## Author contributions

FA conceived the project. FA and JK supervised this study. ALe and ALi sampled the sequenced palms. SS performed the experiments. EC and FA designed the sequence capture. HT, JK, JO and PF performed the analyses of the genomic sequence. HT, JK, FA and PF wrote the manuscript. TB, JK and FA acquired funding. All authors read and approved the final manuscript. JK and FA contributed equally to this work.

## Data and materials availability

Raw reads data are available under the EBI Project PRJEB76653

## Supplementary

### Supplementary notes : Detailed target capture protocol

For each individual, 100 ng of total DNA are sheared using the NEBNext Ultra II FS DNA Modul (New England Biolabs, Ipswich, MA, USA, Part # E7810) by mixing 3.5 µL of NEBNext Ultra II FS Reaction Buffer and 1 µL of NEBNext Ultra II FS Enzyme Mix and an incubation step of 10 min at 37°C. A clean-up step is performed with 1 x volume of Agencourt AMPure XP magnetic beads (Beckman Coulter, Indianapolis, IN, USA, Part # A63881). Fragmented DNA is ligated with 4 pmol of PE-P5 and MPE-P7 adapters. Each PE-P5 and each MPE-P7 adapter carries the same specific hexamer barcode (Rohland & Reich, 2012). Reactions are conducted in 50 µl final volume with 2 unit of T4 DNA ligase (Invitrogen, Thermo Fischer Scientific, Part # 15224025) for 4 hours at 22 °C followed by a heat inactivation step at 65°C for 10 minutes. A clean-up step is performed with 1 x volume of Agencourt AMPure XP magnetic beads. For each individual sample, a pre-hybridization PCR is performed using the KAPA® HiFi HotStart ReadyMix PCR Kit (Roche Diagnostics Indianapolis, IN, USA, Part # KK2602) with 200 nM PreHyb-PE_F (CTTTCCCTACACGACGCTCTTC) and 200 nM PreHyb-MPE_R (TGACTGGAGTTCAGACGTGTG) primers in a final volume of 100µl. Thermocycling parameters were 2 minutes at 98°C, followed by 12 cycles of 20 seconds at 98°C; 45 seconds at 55°C and 30 seconds at 72°C, with a final elongation of 10 minutes at 72°C. A clean-up step is performed with 1 x volume Agencourt AMPure XP magnetic beads. Amplified DNA is individually controlled (sizing and estimation of the concentration) by electrophoresis on a AATI Fragment Analyzer™ (Advanced Analytical Technologies, Ankeny, IA, USA) device with the DNF-474 High Sensitivity Fragment Analysis Kit. Until 24 samples (corresponding to 24 hexamer barcodes on the PE-P5 and PE-P7 adapter) are pooled. A clean-up step is performed with 1 x volume of Agencourt AMPure XP magnetic beads.

Enrichment and capture by hybridization followed the Standard Protocol, with a hybridization and washing temperature of 65°C, following the indications of the User Manual of the myBaits Hybridization Capture for Targeted NGS, Custom DNA-Seq (protocol version 4.01 (http://www.mycroarray.com)). An on-beads PCR amplification is undertaken to enrich library fragments, extend the adapter sequence and incorporate an index to the P7 adapter. The PCR reaction is using the KAPA® HiFi HotStart ReadyMix PCR Kit (KAPABiosystems, Boston, MA, Part # KR0370) in a final volume of 50 µl with:

- 15 pmol of SOL-PE-PCR_F primer (Mascher M *et al*., 2013) aatgatacggcgaccaccgagatctacactctttccctacacgacgctcttc
- 15 pmol of SOL-MPE-INDX_R indexed primers CAAGCAGAAGACGGCATACGAGATXXXXXXGTGACTGGAGTTCAGACGTGT

This primer carries 6 bases of the official TruSeq Illumina Index. Thermocycling parameters were 2 minutes at 98°C, followed by 18 cycles of 20 seconds at 98°C; 30 seconds at 62°C and 30 seconds at 72°C, with a final elongation of 5 minutes at 72°C. The reaction volume is 50 µL. A clean-up step is performed with 1.3 x volume Agencourt AMPure XP magnetic beads. The elution volume is 20 µL. Indexed libraries are individually controlled (sizing and estimation of the concentration) by electrophoresis on an AATI Fragment Analyzer™ device with the DNF-474 High Sensitivity Fragment Analysis Kit. The different indexed libraries, corresponding to all the captured barcoded DNA samples, are equally mixed. The final pooled library is quantified by qPCR with the KAPA Library Quantification Kit (Part # KK4824).

**Table S1.** Classification of sex-linked genes according to SDpop output and X-Y synonymous divergence (dS).

| Gene category <sup>1</sup> | SDpop posterior score | dS |
| --- | --- | --- |
| High-confidence sex-linked (HCSL) | XY score > 0.8 | > 0.0021 *P. dactylifera*<br>> 0.0011 *K. elegans* |
| Probably sex-linked (PSL) | XY score > 0.8<br>0.5 < XY score < 0.8 | < 0.0021 *P. dactylifera*<br>< 0.0011 *K. elegans* |
| Not sex-linked / Autosomal | Autosomal score > 0.8 |  |
| Inconclusive | Other scores |  |
| NA | No data |  |

<sup>1</sup> only applies to genes with more than 10 individuals genotyped on average; other genes are considered “inconclusive”

**Table S2.** Reads mapping of the samples in the *P. dactylifera* WGS dataset.

| Sample name | Reads (R1 + R2) | Mapped properly paired | Discarded<sup>1</sup> |
| --- | --- | --- | --- |
| SRR10121061 | 69906798 | 93.41% | Yes |
| SRR10121067 | 79422680 | 94.90% | Yes |
| SRR10121077 | 94044556 | 95.49% | No |
| SRR10121086 | 70564142 | 94.67% | Yes |
| SRR10121091 | 76352168 | 92.67% | No |
| SRR10121093 | 99377344 | 93.34% | No |
| SRR10121095 | 94123466 | 95.07% | No |
| SRR10121106 | 94660272 | 96.39% | No |
| SRR10121107 | 61350290 | 92.63% | No |
| SRR10121111 | 81614162 | 96.86% | No |
| SRR10121117 | 74139608 | 96.28% | Yes |
| SRR10121120 | 87950120 | 93.51% | No |
| SRR10121126 | 78984780 | 95.37% | No |
| SRR10121130 | 60753842 | 92.48% | No |
| SRR10121134 | 92117304 | 94.42% | No |
| SRR10121138 | 89963210 | 93.65% | No |
| SRR10121146 | 88845434 | 93.90% | No |
| SRR10121160 | 69178418 | 93.86% | No |
| SRR10121164 | 77056038 | 94.57% | No |
| SRR10121166 | 88559374 | 93.80% | No |
| SRR10121171 | 88773146 | 94.72% | No |
| SRR10121175 | 82661366 | 94.40% | No |
| SRR10121177 | 80185434 | 93.98% | No |
| SRR10121178 | 69218370 | 93.77% | No |
| SRR10121179 | 81745170 | 94.27% | Yes |
| SRR10121180 | 77771724 | 92.10% | No |
| SRR10121181 | 85821024 | 93.58% | No |
| SRR10121182 | 110764132 | 94.73% | No |
| SRR10121183 | 98825142 | 94.39% | No |
| SRR10121184 | 77905032 | 94.70% | No |
| SRR10121185 | 81762922 | 94.44% | No |
| SRR10121186 | 96158186 | 94.68% | Yes |

<sup>1</sup> Related individuals (see Figure S2-A) that were discarded from the analysis are indicated by a “Yes”.

**Table S3.** Reads mapping and relatedness of the target capture sequences of SexPhoenix (SP) and PalmExons (PE) datasets.

| Sample | Species | Reads (R1 + R2) |  | Mapped properly paired |  | discarded<sup>1</sup> |
| --- | --- | --- | --- | --- | --- | --- |
|  |  | SP | PE | SP | PE |  |
| Pz187-F | *P. dactylifera* | 4377416 | 18632428 | 99.67% | 99.84% | No |
| Pz188-F | *P. dactylifera* | 5008768 | 18696304 | 99.61% | 99.85% | No |
| Pz189-F | *P. dactylifera* | 2483626 | 9135350 | 99.49% | 99.75% | No |
| Pz190-F | *P. dactylifera* | 4385870 | 17747354 | 99.65% | 99.83% | No |
| Pz191-F | *P. dactylifera* | 2939824 | 10817454 | 99.62% | 99.76% | No |
| Pz192-F | *P. dactylifera* | 3127450 | 10421980 | 99.47% | 99.41% | No |
| Pz193-F | *P. dactylifera* | 1985378 | 6219258 | 99.51% | 99.76% | No |
| Pz194-F | *P. dactylifera* | 3061202 | 11576270 | 99.54% | 99.63% | No |
| Pz195-F | *P. dactylifera* | 2594946 | 9306702 | 99.53% | 99.63% | No |
| Pz196-M | *P. dactylifera* | 2627820 | 10596866 | 99.47% | 99.69% | No |
| Pz197-M | *P. dactylifera* | 3694942 | 16134226 | 99.51% | 99.76% | Yes |
| Pz198-M | *P. dactylifera* | 2281358 | 9850922 | 98.30% | 96.05% | No |
| Pz199-M | *P. dactylifera* | 2078630 | 9343366 | 99.49% | 99.77% | No |
| Pz200-M | *P. dactylifera* | 4513112 | 22881886 | 99.41% | 99.29% | No |
| Pz201-M | *P. dactylifera* | 2016912 | 11571982 | 99.44% | 99.73% | No |
| Pz202-M | *P. dactylifera* | 1772086 | 7567128 | 99.45% | 99.70% | No |
| Pz203-M | *P. dactylifera* | 1941712 | 10554778 | 99.45% | 99.69% | No |
| Pz204-M | *P. dactylifera* | 3704796 | 20085258 | 99.55% | 99.83% | No |
| Pz205-M | *P. dactylifera* | 3084618 | 14996024 | 99.57% | 99.79% | No |
| Pz021-F | *K. elegans* | 666270 | 16383618 | 84.20% | 99.24% | No |
| Pz022-F | *K. elegans* | 752908 | 17161294 | 83.45% | 99.21% | No |
| Pz023-F | *K. elegans* | 771894 | 17218782 | 82.95% | 99.17% | No |
| Pz024-F | *K. elegans* | 738372 | 15364196 | 83.78% | 99.09% | No |
| Pz025-F | *K. elegans* | 874352 | 19668318 | 83.90% | 99.32% | No |
| Pz026-F | *K. elegans* | 603370 | 13660796 | 83.50% | 99.17% | No |
| Pz027-F | *K. elegans* | 576916 | 13522682 | 84.68% | 99.16% | No |
| Pz028-F | *K. elegans* | 752862 | 18841052 | 85.51% | 99.24% | No |
| Pz029-F | *K. elegans* | 926680 | 24343416 | 85.12% | 99.40% | No |
| Pz030-F | *K. elegans* | 733968 | 17460034 | 84.45% | 99.28% | No |
| Pz031-M | *K. elegans* | 789534 | 18569286 | 85.17% | 99.28% | No |
| Pz032-M | *K. elegans* | 439660 | 9782804 | 83.12% | 99.23% | No |
| Pz033-M | *K. elegans* | 677530 | 15510204 | 84.48% | 99.42% | No |
| Pz034-M | *K. elegans* | 500752 | 11488268 | 83.08% | 98.93% | No |
| Pz035-M | *K. elegans* | 709868 | 17164758 | 85.20% | 99.27% | No |
| Pz036-M | *K. elegans* | 575470 | 12718802 | 83.73% | 99.17% | No |
| Pz037-M | *K. elegans* | 623986 | 13381650 | 85.05% | 98.97% | No |
| Pz038-M | *K. elegans* | 691268 | 15951318 | 83.54% | 99.24% | No |
| Pz039-M | *K. elegans* | 942786 | 21623126 | 84.48% | 99.29% | No |
| Pz040-M | *K. elegans* | 756512 | 18298996 | 85.16% | 99.39% | No |
| Pz269-F | *K. elegans* | 578186 | 12884964 | 83.44% | 98.83% | No |
| Pz270-H | *K. elegans* | 595258 | 14863394 | 83.04% | 99.06% | Yes |
| Pz271-M | *K. elegans* | 755654 | 19055042 | 84.54% | 99.39% | No |
| Pz272-F | *K. elegans* | 508120 | 11612810 | 81.86% | 99.03% | No |
| Pz273-F | *K. elegans* | 1014112 | 26035728 | 82.67% | 99.10% | No |
| Pz274-F | *K. elegans* | 607980 | 12829890 | 82.60% | 99.20% | No |

<sup>1</sup> Related individuals that were discarded from the analysis are indicated by a “Yes”.

**Table S4.** Read depth and coverage of the DPV01 genome protein-coding genes by our different datasets.

|  | P. dactylifera |  |  | K. elegans |  |
| --- | --- | --- | --- | --- | --- |
|  | WGS<sup>1</sup> | SP<sup>2</sup> | PE<sup>3</sup> | SP<sup>2</sup> | PE<sup>3</sup> |
| DPV01 genes covered (%)<sup>4</sup> | 97.61 | 1.6 | 77.06 | 2.54 | 58.31 |
| Median covered length (%)<sup>5</sup> | 99.65 | 74.80 | 48.30 | 53.80 | 43.30 |
| [1<sup>st</sup> - 3<sup>rd</sup> quartile ] (%) | [92.25 - 1] | [48.29 – 94.50] | [33.15 – 66.74] | [23.23 – 87.14] | [29.15 – 59.94] |
| Median depth per base (females)<sup>6</sup> | 11.03 | 39.01 | 22.43 | 19.89 | 32.50 |
| [1<sup>st</sup> - 3<sup>rd</sup> quartile ] | [10.07 – 11.74] | [11.62 – 93.24] | [8.51 – 48.42] | [1.86 – 82.25] | [9.50 – 87.91] |
| Median depth per base (males)<sup>6</sup> | 11.76 | 36.07 | 19.36 | 18.93 | 31.18 |
| [1<sup>st</sup> - 3<sup>rd</sup> quartile ] | [10.85 – 12.40] | [15.01 – 80.80] | [7.70 – 40.75] | [1.88 – 78.84] | [9.10 – 84.67] |
| Number of variant positions analyzed | 466719 | 4693 | 180035 | 8395 | 161384 |

<sup>1</sup> Whole Genome Sequencing (WGS)
<sup>2</sup> SexPhoenix capture kit (SP)
<sup>3</sup> PalmExons capture kit (PE)
<sup>4</sup> Percentage of protein coding genes covered from the DPV01 P. dactylifera genome annotation.
<sup>5</sup> Median, first and third quartile of the proportion of the total gene length covered over 10X for WGS and over 7X for SP and PE.
<sup>6</sup> Median, first and third quartile of the mean depth of coverage per base per gene.

**Table S5.**
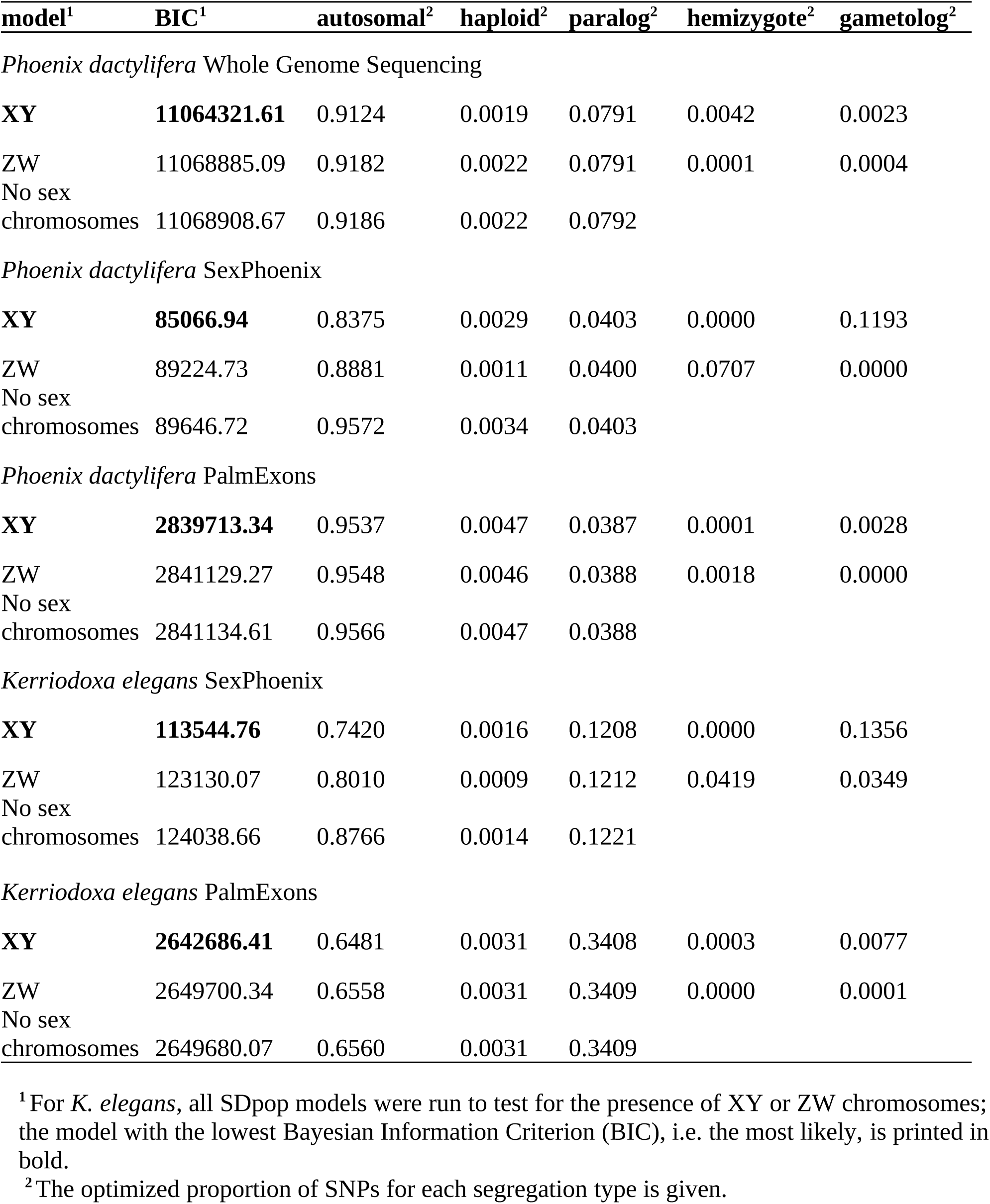
Summary of SDpop model analysis.

**Table S6:** Summary of the number of genes in Whole Genome Sequencing (WGS) and <u>PalmExons capture (PE) datasets and position of sex-linked genes.</u>.

| Number of genes | Pd<sup>1</sup> WGS | Pd PE | Ke<sup>1</sup> PE |
| --- | --- | --- | --- |
| Total (in analysis) | 25 100 | 19 889 | 15 051 |
| High-confidence or probably sex-linked<sup>3</sup> | 87 | 68 | 89 |
| *P. dactylifera* PDMBC4 genome |  |  |  |
| Chr12:9.4-12.4Mb<sup>4</sup> | 22 | 12 | 16 |
| *P. dactylifera* PDMBC4 genome |  |  |  |
| unplaced scaffold<sup>4</sup> | 39 | 25 | 36 |
| *E. guineensis*<sup>5</sup> EGPMv6 genome |  |  |  |
| Chr10:43-46.1Mb<sup>4</sup> | 52 | 29 | 55 |
| *E. guineensis* EGPMv6 genome |  |  |  |
| unplaced scaffold<sup>4</sup> | 0 | 0 | 2 |

<sup>1</sup> *Phoenix dactylifera*
<sup>2</sup> *Kerriodoxa elegans*
<sup>3</sup> see Material and Methods
<sup>4</sup> Among High-confidence or probably sex-linked
<sup>5</sup> *Elaeis guineensis*

**Table S7.**
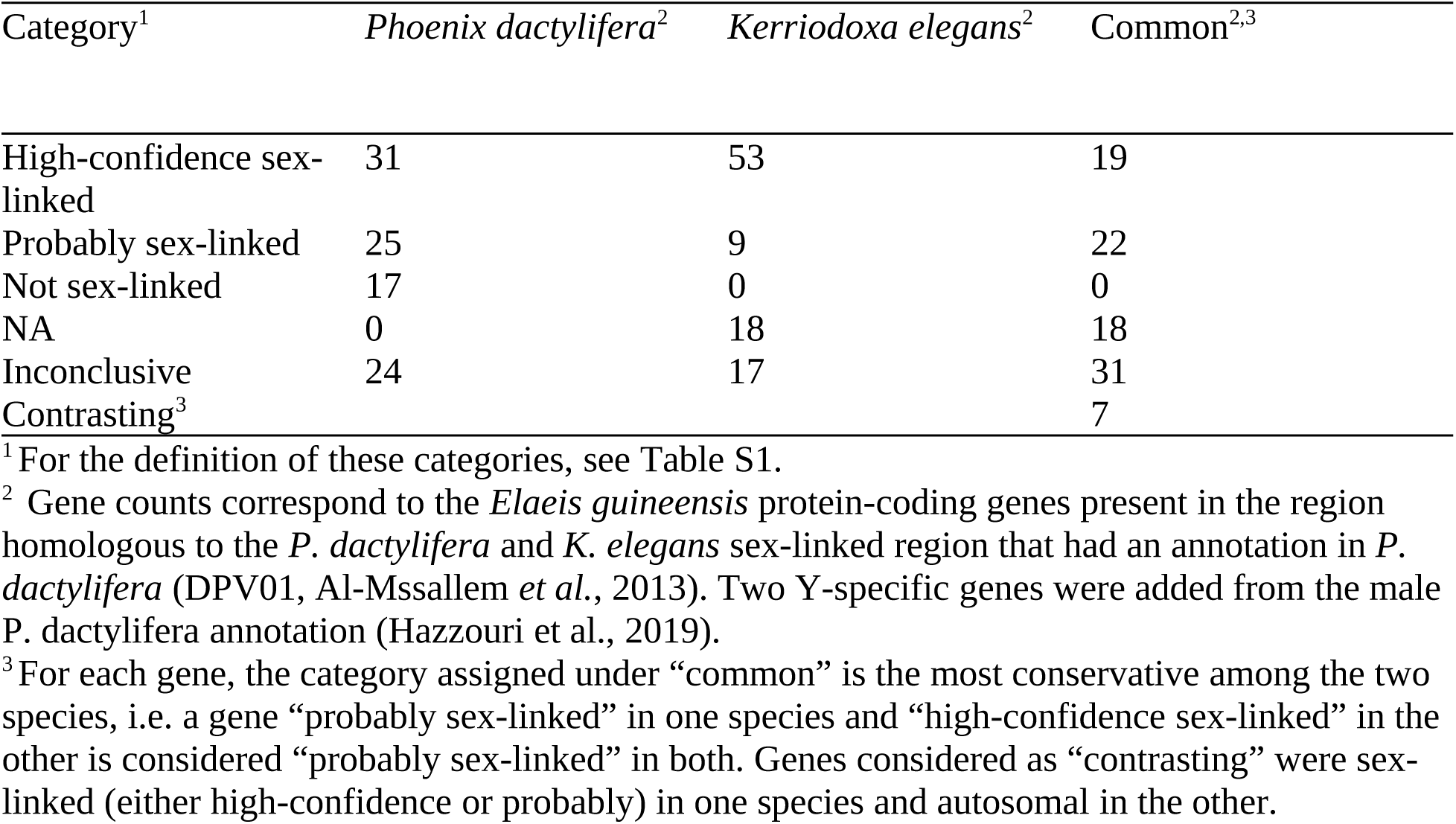
Classification of protein-coding genes in the region of *E. guineensis* homologous with the sex-linked region in *P. dactylifera* and *K. elegans*.

**Table S8.** XY sequences for which gene conversion was detected.

| Gene | XY paired sequences<sup>1</sup> | Simulated p-value<sup>2</sup> | KA p-value<sup>2</sup> | start position<sup>3</sup> | end position<sup>3</sup> |
| --- | --- | --- | --- | --- | --- |
| LOC103695972 | *P. dactylifera* WGS<sup>4</sup> | 0.01 | 0.03 | 1013 | 1214 |
| LOC103695972 | *P. dactylifera* SP<sup>5</sup> | 0.01 | 0.03 | 1013 | 1214 |
| LOC103698366 | *P. dactylifera* WGS | 0.02 | 0.09 | 73 | 716 |
| LOC103698366 | *P. dactylifera* SP | 0.01 | 0.05 | 73 | 716 |
| LOC103716262 | *K. elegans* SP | 0.01 | 0.08 | 250 | 699 |
| LOC103723089 | *K. elegans* SP | 0.02 | 0.14 | 448 | 1061 |
| LOC103724019 | *P. dactylifera* WGS | 0.02 | 0.08 | 3299 | 4064 |
| LOC103724019 | *P. dactylifera* SP | 0.03 | 0.11 | 3299 | 4064 |
| LOC103723285 | *K. elegans* SP | 0.00 | 0.02 | 686 | 824 |

<sup>1</sup> Dataset of gene sequences where gene conversion was detected between the X and Y copies.
<sup>2</sup> KA p-values are more conservative and less accurate than simulated p-values. Only one was required to be below 0.05 to be included.
<sup>3</sup> Start and end position of the alignment which underwent gene conversion.
<sup>4</sup> Whole Genome Sequencing (WGS) data.
<sup>5</sup> SexPhoenix (SP) target capture data.

**Table S9.** Synonymous divergence (dS) between *Elaeis guineensis* and *Phoenix dactylifera* for ten random autosomal single copy orthologs.

| *Elaeis guineensis*<br>gene | *Phoenix dactylifera*<br>gene | Gene function | dS |
| --- | --- | --- | --- |
| LOC105059023 | LOC103698418 | DNA repair protein RAD51 homolog 3 | 0.18 |
| LOC105056566 | LOC103703289 | thyroid adenoma-associated protein homolog | 0.12 |
| LOC105056732 | LOC103705344 | pentatricopeptide repeat-containing protein OGR1 | 0.23 |
| LOC105057008 | LOC103706629 | nudix hydrolase 9 | 0.15 |
| LOC105036130 | LOC103708307 | FT-interacting protein 7 | 0.30 |
| LOC105049520 | LOC103711984 | chloride channel protein CLC-e | 0.12 |
| LOC105052889 | LOC103713882 | tetratricopeptide repeat protein SKI3 | 0.10 |
| LOC105047177 | LOC103714988 | anaphase-promoting complex subunit 10 | 0.15 |
| LOC105057153 | LOC103720210 | uncharacterized | 0.22 |
| LOC105057944 | LOC103720756 | uncharacterized | 0.06 |

**Figure S1.**
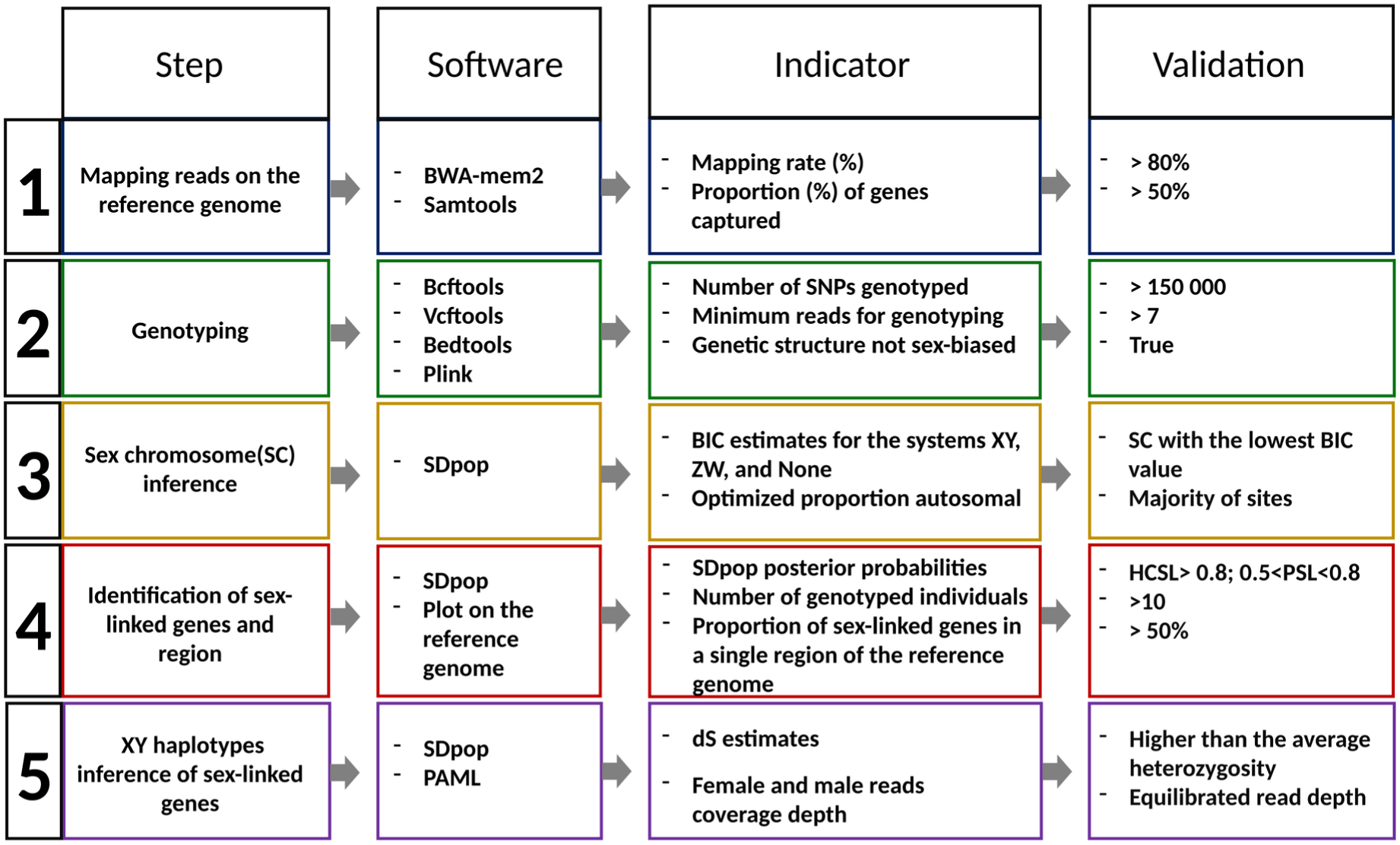
Strategy summary. Here are the main steps of our analysis with the softwares, their variables and threshold values used for step validation.

**Figure S2.**
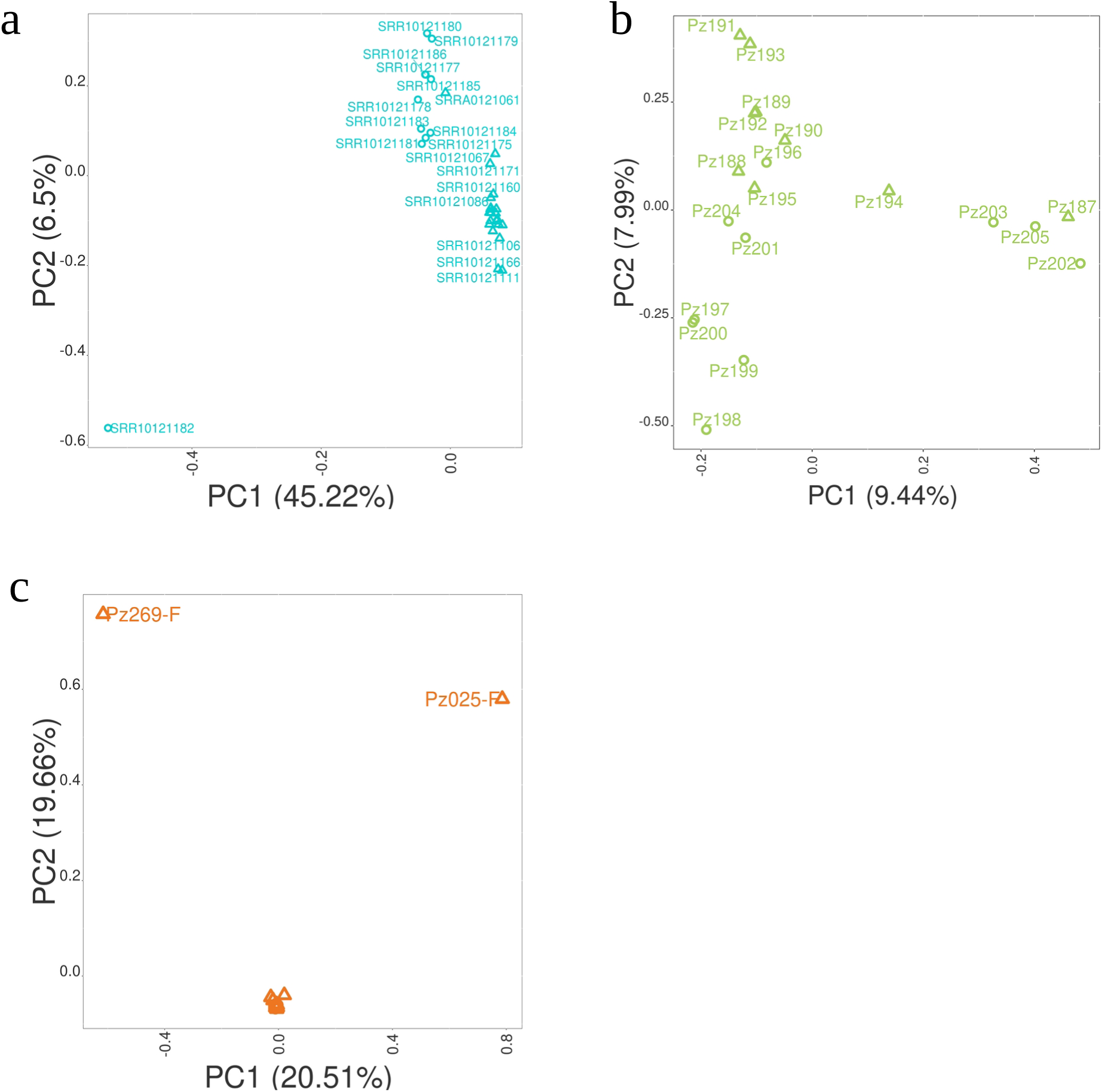
PCA of the genetic variation of studied populations. Females are represented by triangles, males by circles, one ambiguous *K. elegans* individual by a square. **(a)** PCA of the genetic variation of the *P. dactylifera* whole genome individuals. **(b)** PCA of the genetic variation captured with PalmExons kit among the *P. dactylifera* individuals. **(c)** PCA of the genetic variation captured with PalmExons kit among the *K. elegans* individuals.

**Figure S3.**
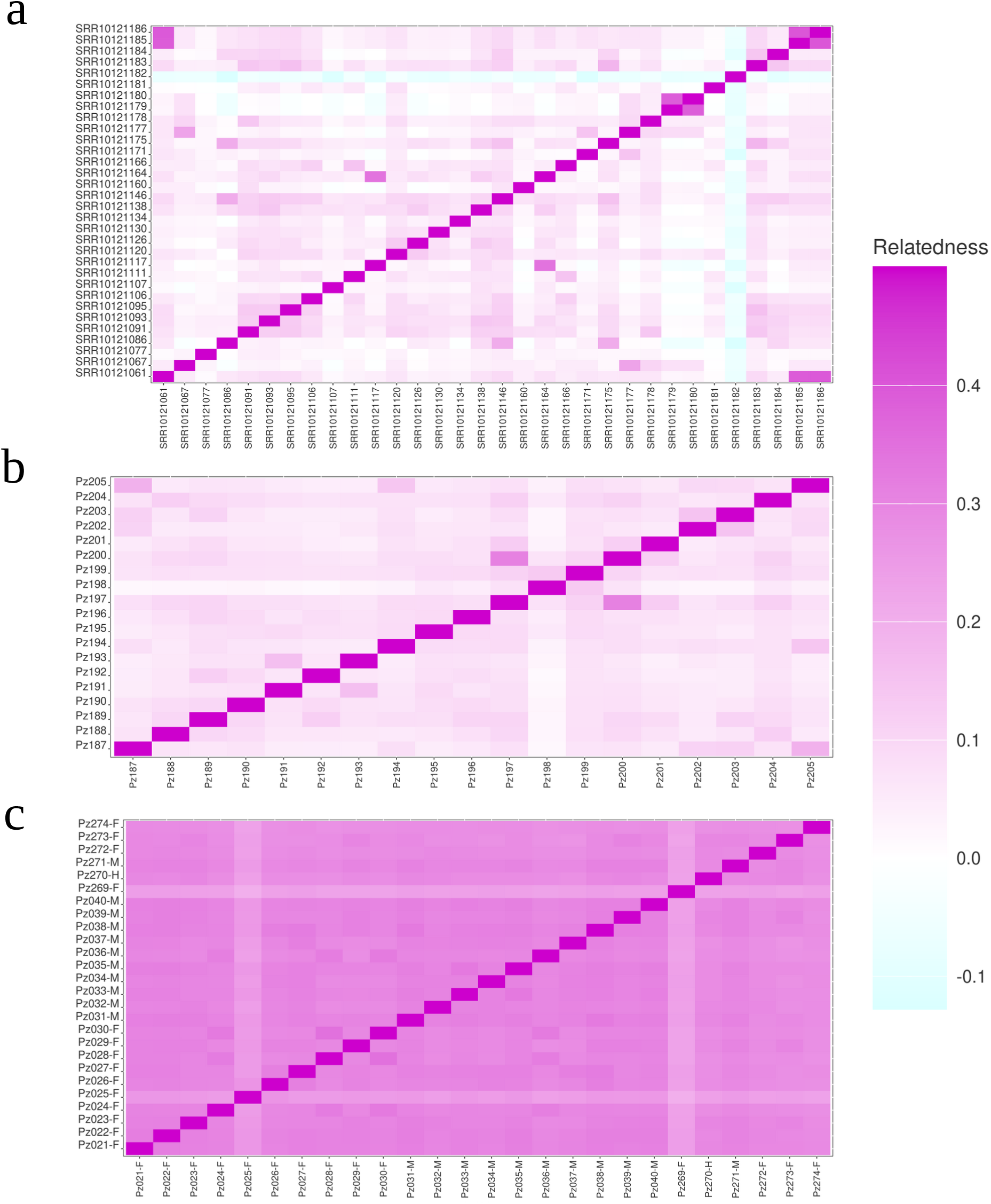
Heatmap of estimated relatedness between individuals. A relatedness of 0 is represented by a white square and indicates unrelated individuals from the same population. A relatedness of 0.5 (the maximum value) is represented with a magenta square and is characteristic of clones. A cyan square indicates negative relatedness as one would expect between different populations. Gradients between these colors represent intermediate values. **(a)** Heatmap of the relatedness between *P. dactylifera* whole genome individuals. **(b)** Heatmap of the relatedness between *P. dactylifera* individuals in the data captured with PalmExons kit. **(c)** Heatmap of the relatedness between *K. elegans* individuals in the data captured with PalmExons kit.

**Figure S4.**
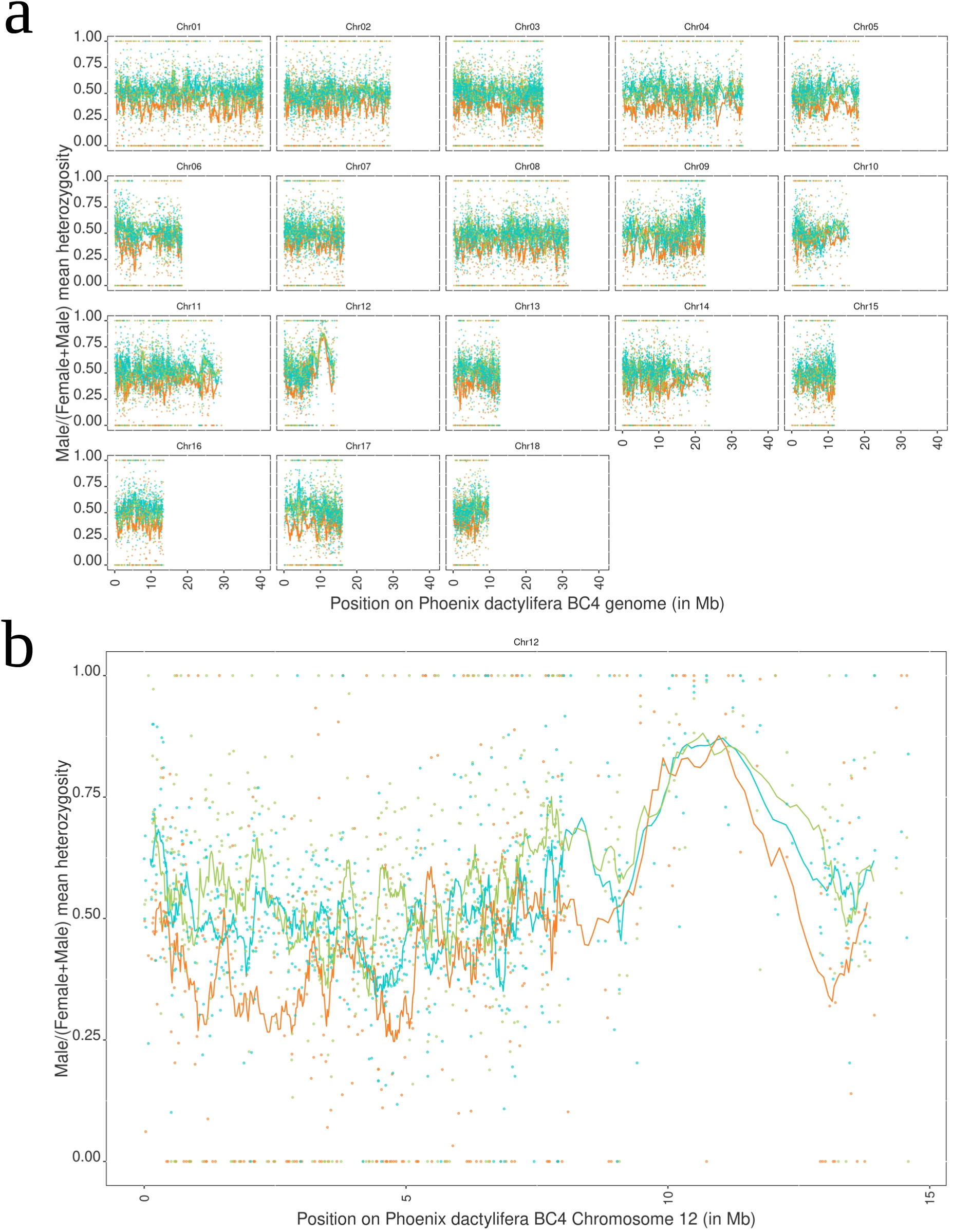
**Male mean heterozygosity proportion per gene mapped on PDMBC4 genome of *P. dactylifera.*** Male (M) and female (F) mean heterozygosity rate per gene on all genotyped positions were calculated. Genes without heterozygosity were removed. Then, the ratio of male mean heterozygosity over female and male mean heterozygosity was calculated : M/(M+F). **(a)** Male mean heterozygosity proportion per gene is represented on the chromosomes of the *Phoenix dactylifera* PDMBC4 genome. **(b)** Male mean heterozygosity proportion per gene is represented on the 12th chromosome of the PDMBC4 genome. Each line represents the rolling mean over 15 genes colored according to the species and dataset : *P. dactylifera* Whole Genome Sequencing (WGS; cyan), PalmExons capture sequence for *P. dactylifera* (green) and *K. elegans* (orange).

**Figure S5.**
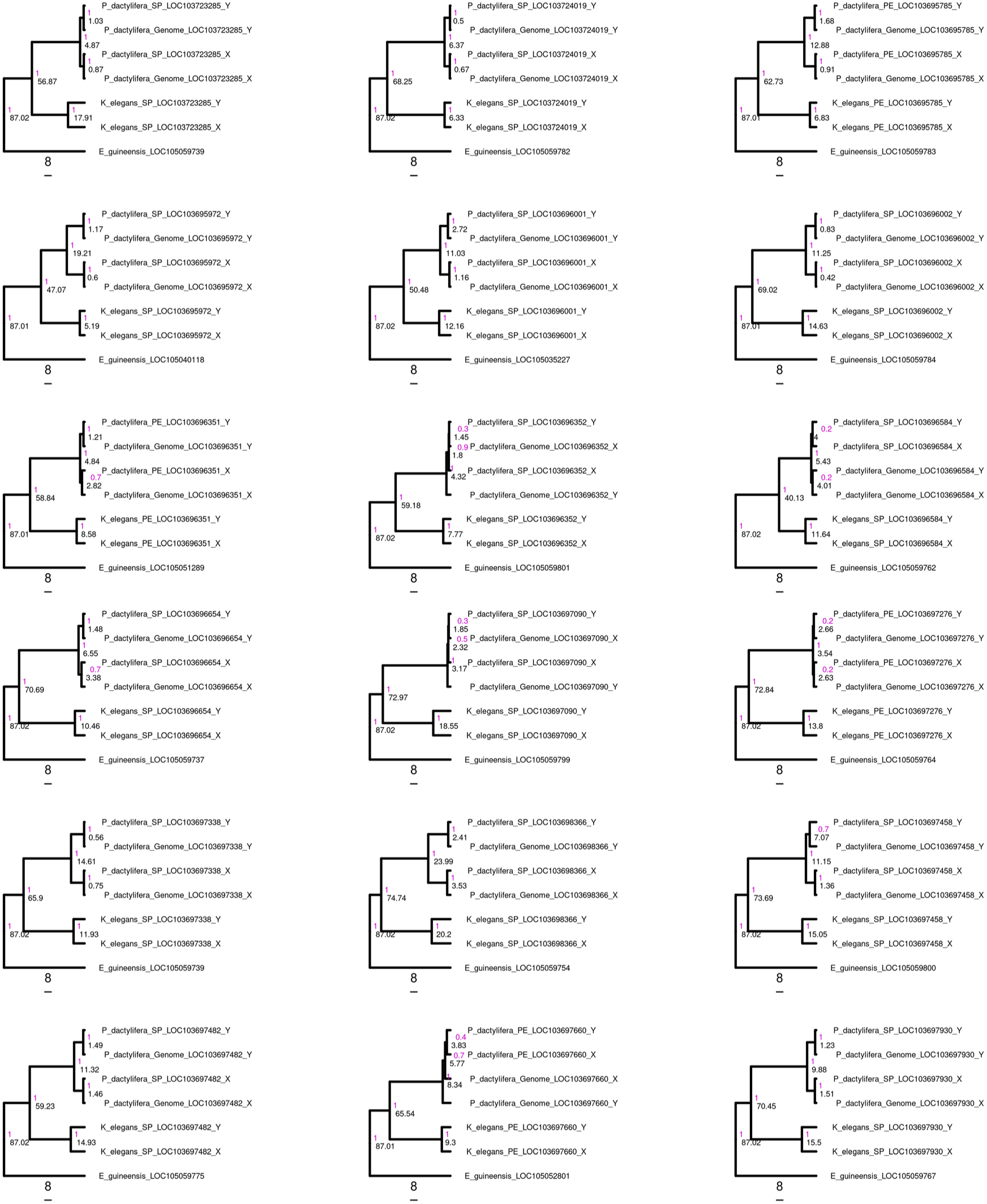
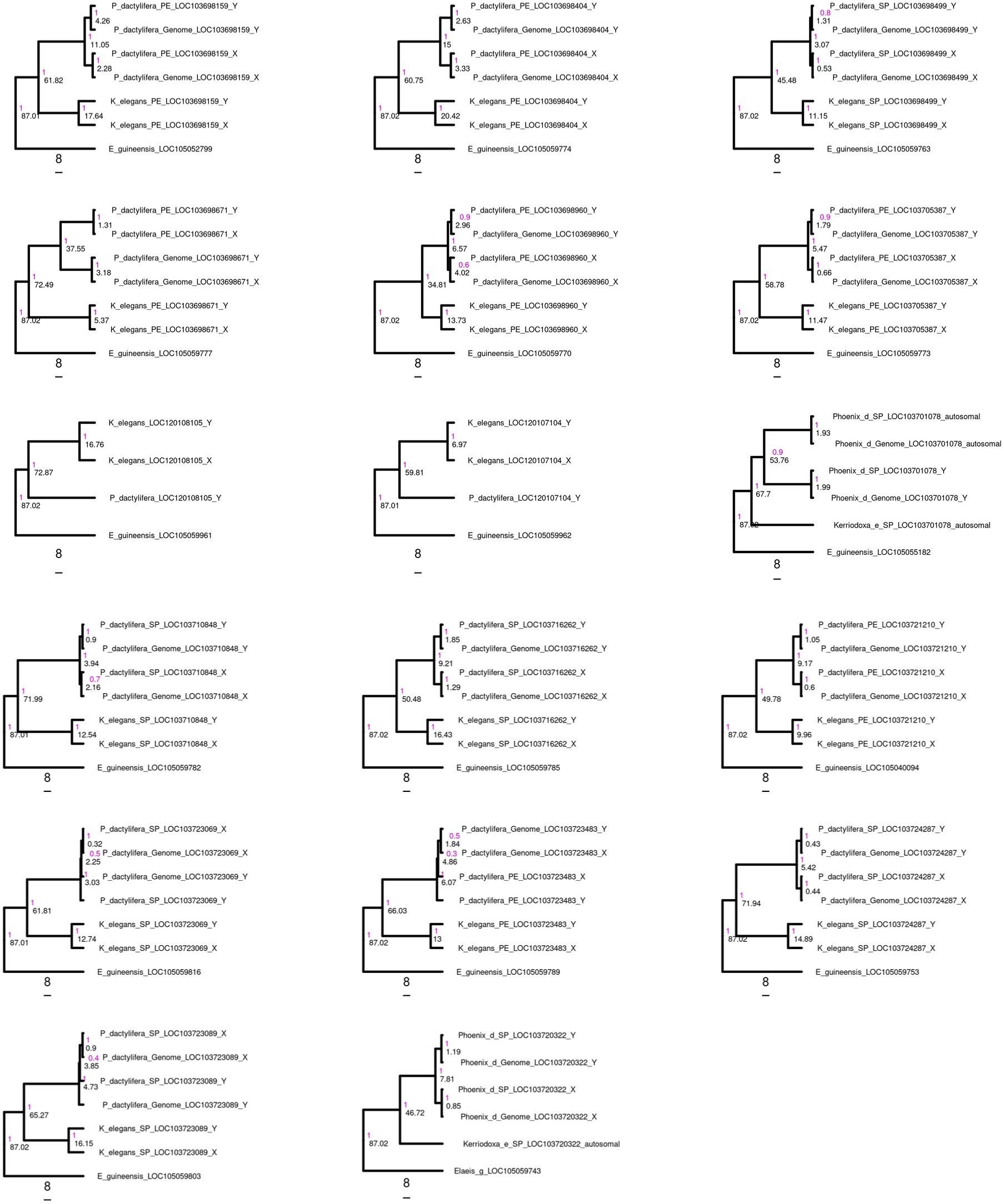
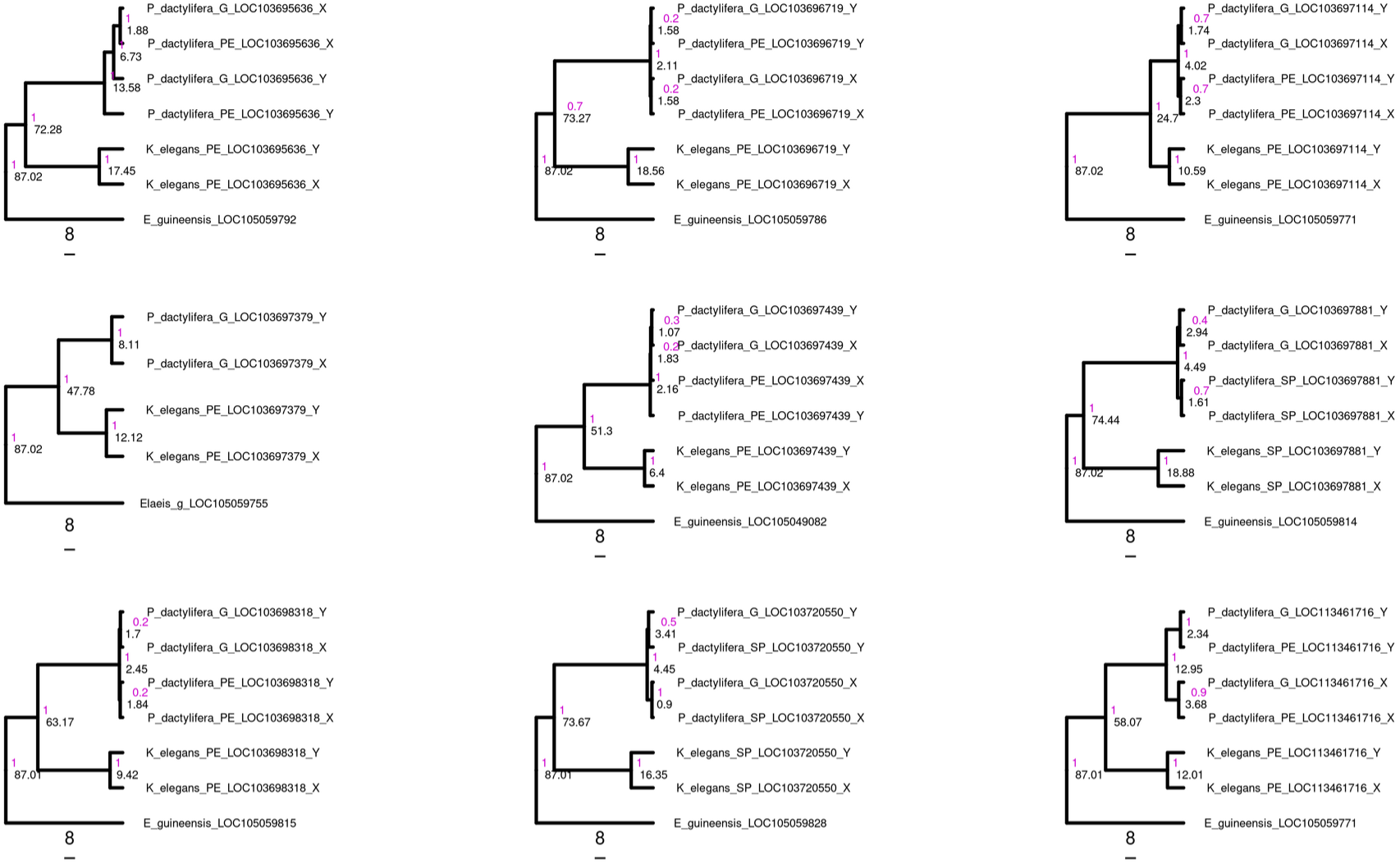
Phylogenies of sex-linked genes. Phylogenies of all sex-linked genes common to *K. elegans* and *P. dactylifera* (high-confidence sex-linked and probably sex-linked, File **S3**) and two known sex-linked genes in *P. dactylifera* : LOGlike (LOC103701078) and guanosine deaminase (LOC103720322). Two Y-specific sequences of *P. dactylifera* were taken from NCBI : GPAT3 (LOC120108105, MH668906), CYP703 (LOC120107104, MH668888). *Elaeis* guineensis root sequences were also taken from NCBI as the blastn top hit of *P. dactylifera* sequence. The node values are the posterior probabilities of the node colored in magenta and the mean age of the node in black. The scales below the trees are in million years.

**Figure S6.**
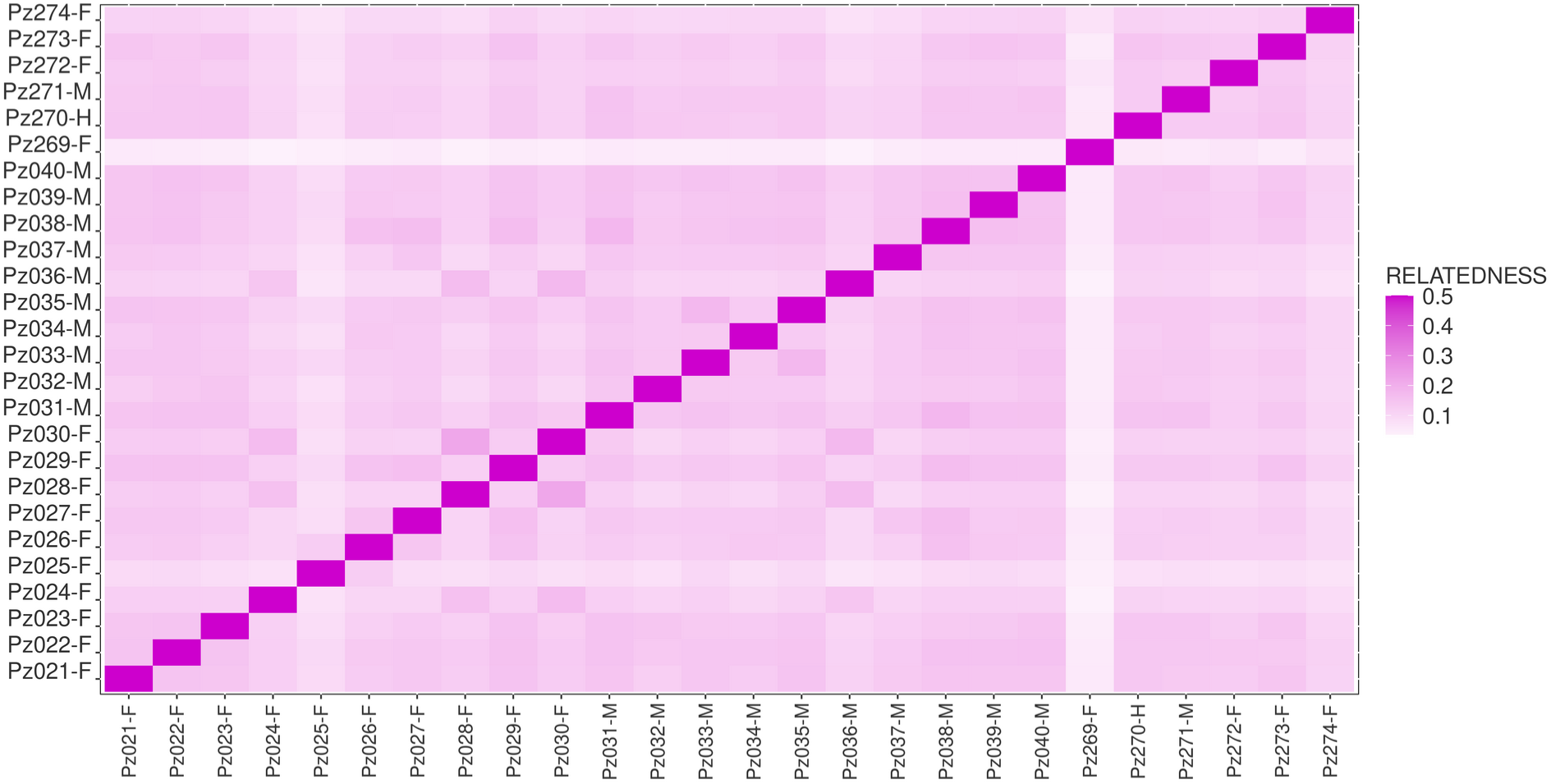
*Kerriodoxa elegans* heatmap of estimated relatedness with no paralogy bias. Genes with high posterior probabilities to be paralogous were removed from the data prior to estimating relatedness. A relatedness of 0 is represented by a white square and indicates unrelated individuals from the same population. A relatedness of 0.5 (the maximum value) is represented with a magenta square and is characteristic of clones. Gradients between white and magenta represent intermediate values.

**Figure S7:**
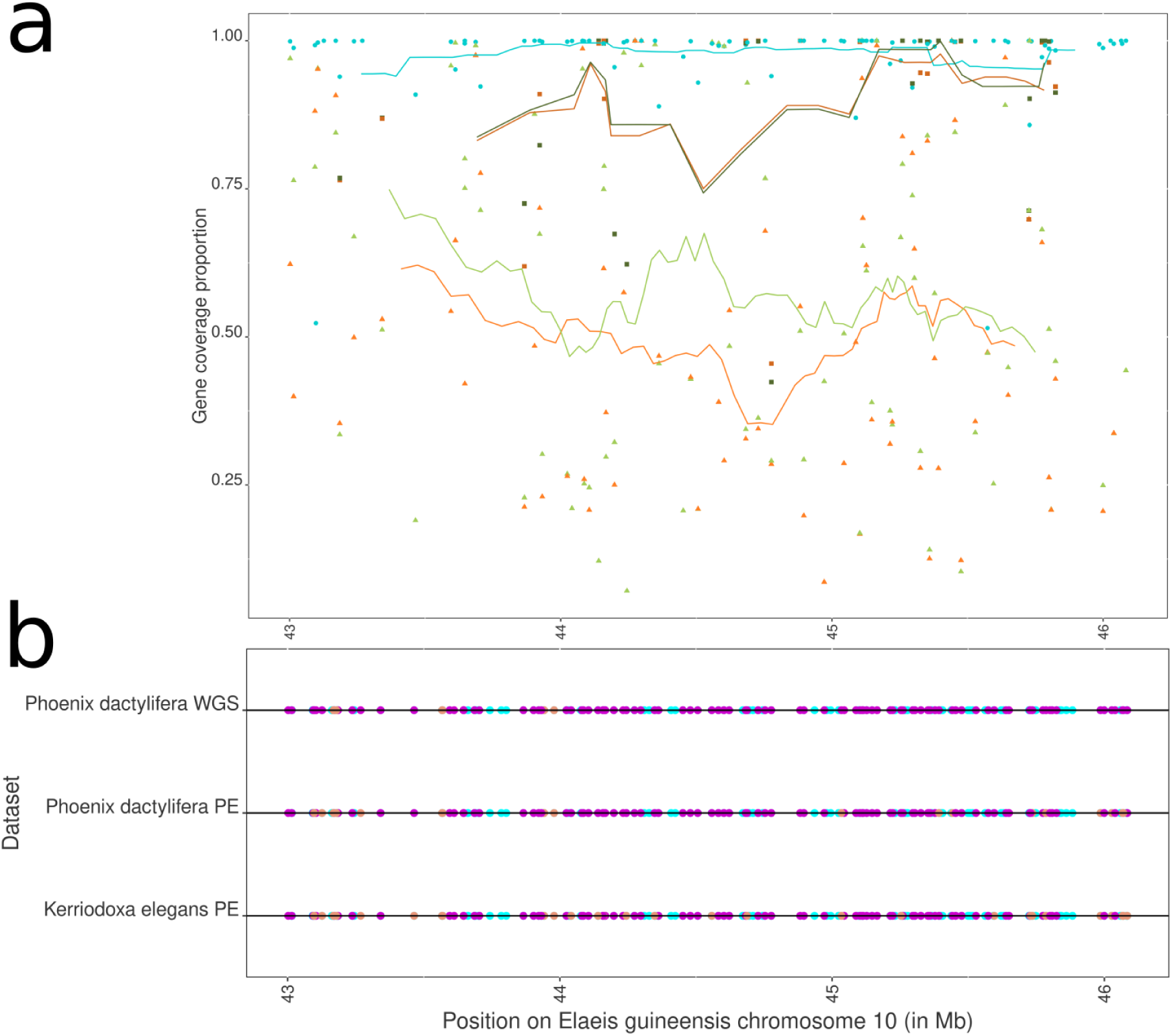
Sex-linked region coverage. **(a)** Proportion of each gene covered (as a fraction of the total length) with at least 7X depth for capture kits and 10X for whole genome sequencing (WGS) in the sex-linked region of *Phoenix dactylifera* and *Kerriodoxa elegans, using the corresponding region* on the *Elaeis guineensis* EGPMv6 genome as a reference. Each point represents a gene for the WGS (circle), the PalmExons sequence capture (triangle) and SexPhoenix sequence capture (square). They are colored in blue (WGS *Phoenix dactylifera*), green (sequence capture *P. dactylifera*), orange (sequence capture *Kerriodoxa elegans*) with lighter colors for PalmExons data and darker for SexPhoenix. **(b)** *P. dactylifera* DPV01 and *E. guineensis* EG5 genome annotations genes positioned via Liftoff on the *E. guineensis* EGPMv6 genome 10^th^ chromosome between 43Mb and 46.1Mb. *P. dactylifera* genes are colored depending on their presence (magenta) or absence (lightpink) in a dataset (WGS or PE) and positions with genes unique to *E. guineensis* are colored in cyan.

**Figure S8.**
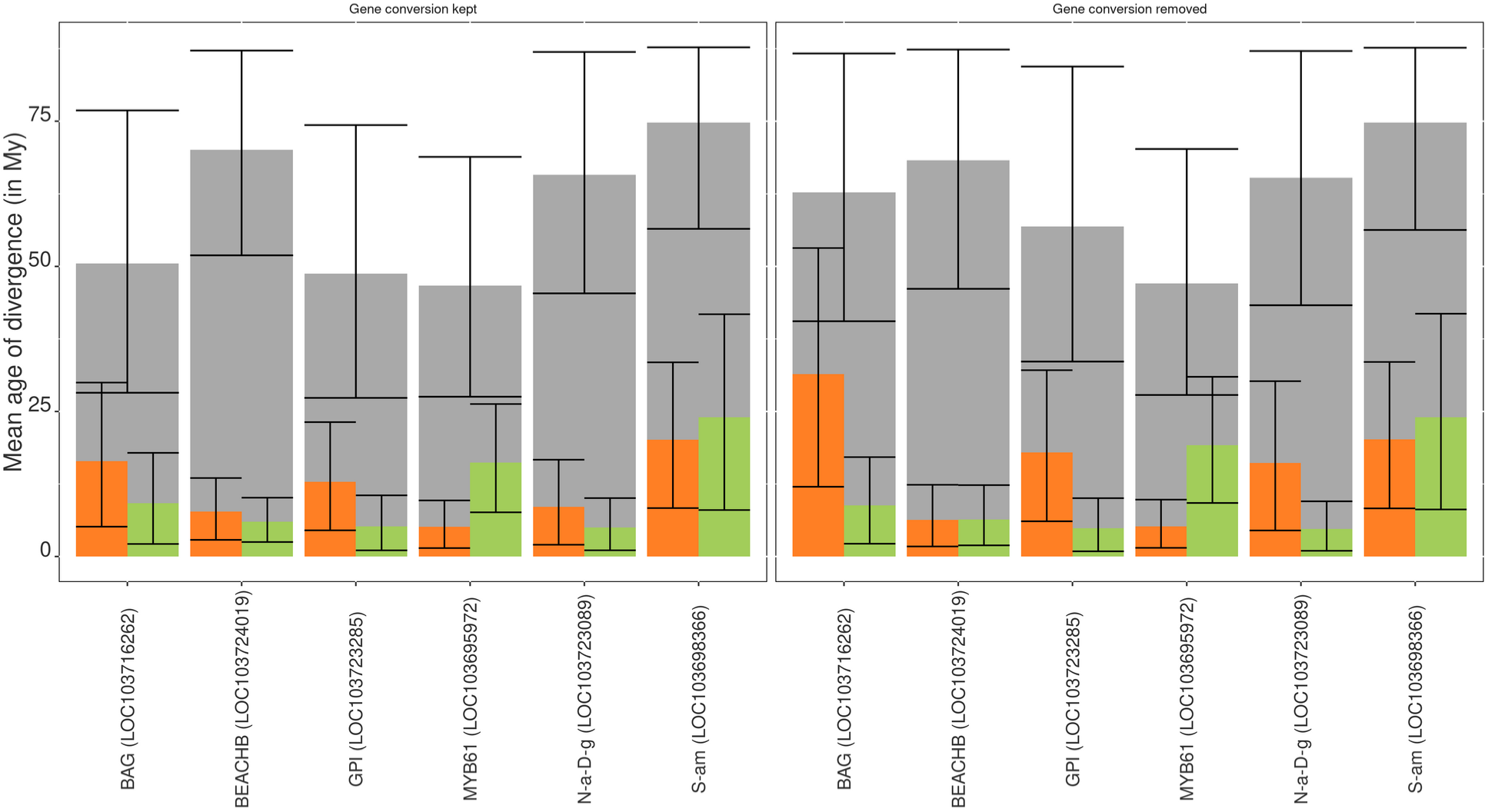
Effect of gene conversion between X and Y copies on node ages. Mean age of divergence of X and Y sequences for common sex-linked genes with possible gene conversion. Gene fragments detected as having undergone gene conversion (see Table S8) were either retained in the analysis (left panel) or removed (right panel). *P. dactylifera* sex-linked sequence divergence estimates are colored in green, estimated for *K. elegans* in orange, and the estimates of the age of the split of the lineages between the two species in grey. Error bars correspond to the 95% probability density of the node age.

**Figure S9.**
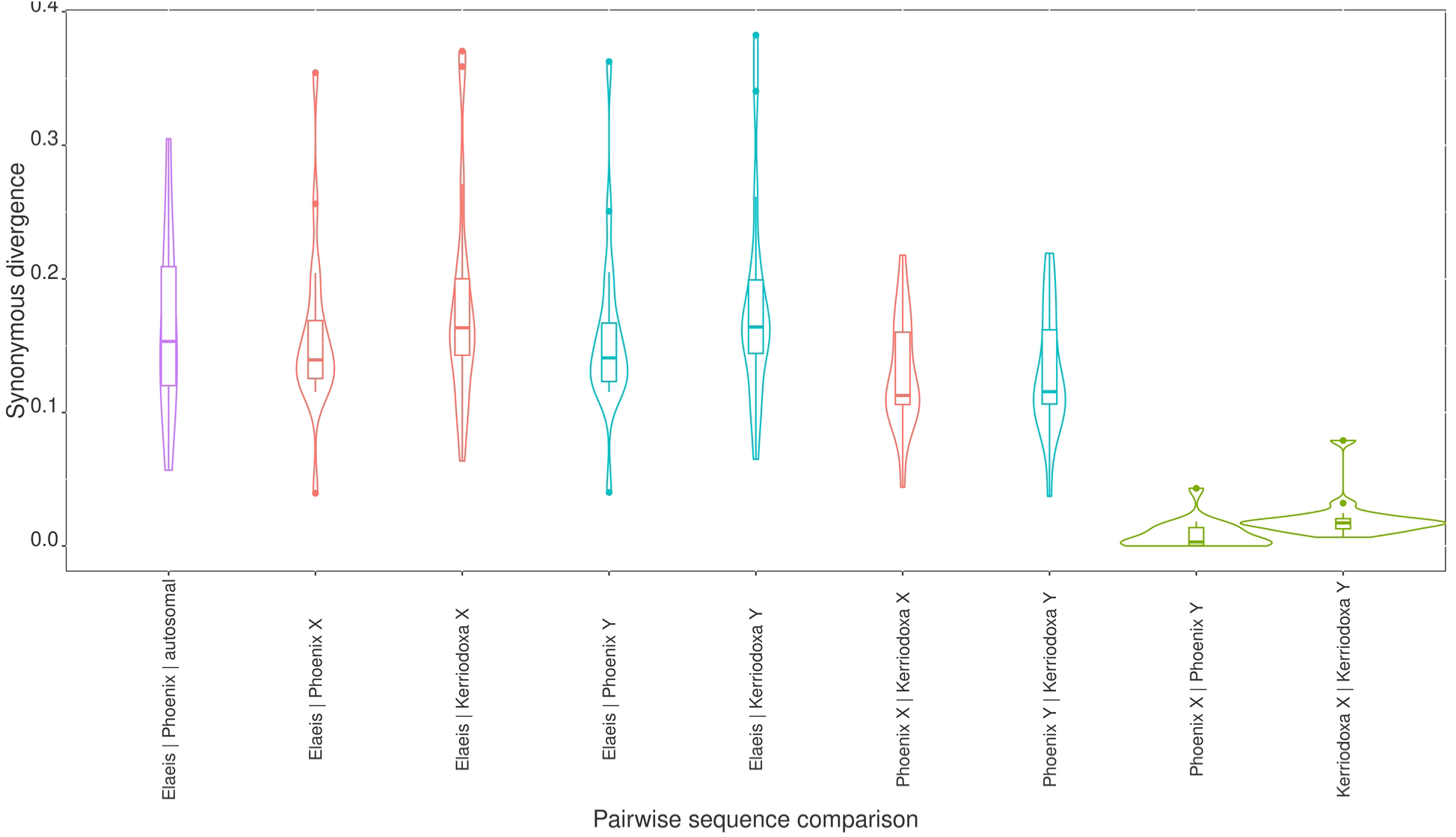
Synonymous divergence of sex-linked and autosomal nuclear genes. Violin plots and boxplots of the synonymous divergence (dS) of the 18 genes inferred as XY in SexPhoenix (SP) and 10 randomly selected autosomal single copy orthologs pertaining to the BUSCO *Liliopsidae* database. dS between X and Y sequences are colored in green, dS between X sequences from SP are colored in red, dS between Y sequences from SP are colored in cyan and dS between autosomal single copy orthologs are colored in purple.

**File S1 (separate file). SexPhoenix target reference sequences available in EBI Project PRJEB76653.**

**File S2 (separate file). PalmExons baits sequences available in EBI Project PRJEB76653.**

**File S3. (separate file). Summary of all genes either inferred as XY in at least one dataset or aligned on the EGPMv6 genome (Ong *et al*., 2020) on chromosome 10 between 43Mb and 46.1Mb.** Each gene identification and annotation is documented as well as their position on the PDMBC4 genome (Hazzouri *et al*., 2019), EG5 annotation (Singh *et al*., 2013) and EGPMv6 (Ong *et al*., 2020). This table contains the mean female and male depth per base as well as the mean female reads ratio, the number of polymorphisms and mean number of individual genotyped per polymorphism as well as the autosomal and XY posterior probability of each of these genes.

**File S4. (separate file). Summary of all genes inferred as X-hemizygous in at least one dataset.** Each sheet contains the X-hemizygous genes of one dataset. Since no X-hemizygous genes were found in any SexPhoenix dataset, no sheet is present in the file for these datasets. Each gene identification and annotation is documented as well as their position on the PDMBC4 genome (Hazzouri *et al*., 2019), EG5 annotation (Singh *et al*., 2013) and EGPMv6 (Ong *et al*., 2020). This table contains the mean female and male depth per base as well as the mean female reads ratio, the number of polymorphisms and mean number of individual genotyped per polymorphism as well as the X-hemizygous posterior probability of each of these genes.

**File S5. (separate file). GO annotation of the DPV01 P. dactylifera genome.** Merged GO annotation of the DPV01 genome through the fantasia deep-learning method with the PDMBC4 P. dactylifera genome GO annotation of genes common to both assemblies’ annotations. Each gene identification is referenced with the GO ID of the annotated GO terms for said gene.

**File S6. (separate file). GO enrichment of the common sex-linked region: *Elaeis guineensis* EGPMv6 genome (Ong et al., 2020) chromosome 10, 43.3Mb to 46.1Mb.** Enriched GO-IDs and terms are summarised. The number of genes annotated for each GO-id in the universe, the number present in the region of interest as well as the expected number are referenced with the estimated p-value of enrichment and the reference universe. SDpop-1X and SDpop-7X are the two gene universes used and are the list of genes present in the SDpop analysis with a respective mean depth per base of 1 and 7.

